# Single-nucleus profiling of social behaviour in a mutualistic reef fish

**DOI:** 10.64898/2026.09.15.751910

**Authors:** Debora Desantis, David J. Wright, Wilfried Haerty, Ashleigh Lister, Iain Macaulay, Celia Schunter

## Abstract

Complex social decision-making in vertebrates relies on a conserved network within the forebrain. However, the precise neural cell populations and gene regulatory networks that orchestrate these sophisticated social behaviours remain largely unresolved. Here, we present the first single-nucleus transcriptome atlas of the telencephalon in the cleaner wrasse *Labroides dimidiatus,* a remarkable model for advanced social cognition. Profiling ∼67,000 telencephalic nuclei, we mapped the neural architecture of the teleost forebrain and identified highly conserved cell populations including a cholinergic-GABAergic neuron population implicated in social behaviours and cells resembling mammalian striatal medium spiny neurons and hippocampal cell types. By comparing distinct behavioural states, we found that socially interacting fish expanded radial glial subclusters, and associative learning co-expression networks, whereas social observers exhibited gene expression signatures for motivation and behavioural preparedness. Ultimately, the resolved telencephalon atlas reveals the evolutionarily conserved cellular and molecular substrates that may govern complex social behaviour across vertebrates.

## INTRODUCTION

Cooperative social behaviours, particularly mutualism, require profound cognitive abilities, demanding individuals to continuously assess risk, reward and dynamic social context (*1*, *2*). While the neural basis of complex behavioural responses is typically studied in mammalian models, the coral reef blue streak cleaner wrasse (*Labroides dimidiatus,* herein ‘cleaners’) exhibits a level of social intelligence and cognitive capability remarkably convergent with primates (*3*, *4*). As a fundamental marine species, cleaners sustain reef biodiversity by engaging in mutualistic interactions to remove ectoparasites and dead tissue from other fishes defined as “clients” (*5–8*). This service not only controls parasite loads (*9*, *10*), but also reduces client stress (*11*) and enhances client cognitive performance (*12*). This mutualism requires sophisticated decision-making strategies: cleaners must constantly balance the temptation to “cheat” by consuming the nutrient-rich client mucus against the risk of aggressive punishment, and they frequently employ tactile stimulation to reconcile from conflicts and secure future interactions (*11*, *13–15*). Consequently, *L. dimidiatus* provides an unprecedent, ecologically relevant model for decoding the evolutionary and cellular neurobiology underlying complex interspecies social cooperation.

In higher vertebrates, complex social decision-making, learning, and memory are orchestrated by a conserved network of forebrain regions, which in the teleosts brain are homologous to the telencephalon (*16*, *17*). Within this network, neuromodulators such as dopamine and serotonin are known to govern the motivation to interact in cleaners (*18*, *19*). Furthermore, bulk transcriptomic studies in cleaners and other fishes have linked interaction behaviours and client recognition to the regulation of glutamate pathways, calcium channel subunits, and Immediate Early Genes (IEGs) such as *c-fos* (*20*, *21*). However, traditional bulk RNA sequencing obscures the functional heterogeneity of the brain (*22*). Because this sequencing method averages signals across diverse tissues, the precise neural cell populations and the dynamic gene regulatory networks that execute these advanced social behaviours remain a critical open question (*23*). To truly understand how the vertebrate brain encodes cooperation, we must deconstruct the social brain at single-cell resolution. Cleaners provide an ideal system for achieving this experimentally, as their behaviour can be easily manipulated and the neural effects of controlled social stimulation can be measured directly.

Here, we resolve the cellular architecture of a complex social (mutualistic) behaviour by integrating single-nuclei RNA sequencing (snRNA-seq) with behavioural profiling of the cleaner wrasse telencephalon (Fig. 1, A to F). We generate the first comprehensive neural cell atlas for a highly cooperative marine vertebrate, enabling cross-species comparisons of telencephalic cell types. By comparing actively socially interacting individuals with social observer fish, we map sociality driven transcriptional dynamics, cellular excitation, and co-expression gene networks to distinct neural cell populations. Given the highly conserved nature of the vertebrate social decision-making network (*17*), we hypothesize that complex interaction behaviours trigger highly specific transcriptional cascades, particularly involving IEGs and glutamatergic plasticity (*20*, *24*) within discrete neuronal subpopulations. Furthermore, we evaluate whether neuromodulatory regulation of cooperation correlates with distinct transcriptomic shifts within dopaminergic and serotonergic pathways (*19*, *25–27*). Ultimately, our findings bridge the gap between neuroanatomical cell architecture and complex ecological function, revealing the fundamental cellular targets that drive vertebrate social behaviour.

**Fig. 1.**
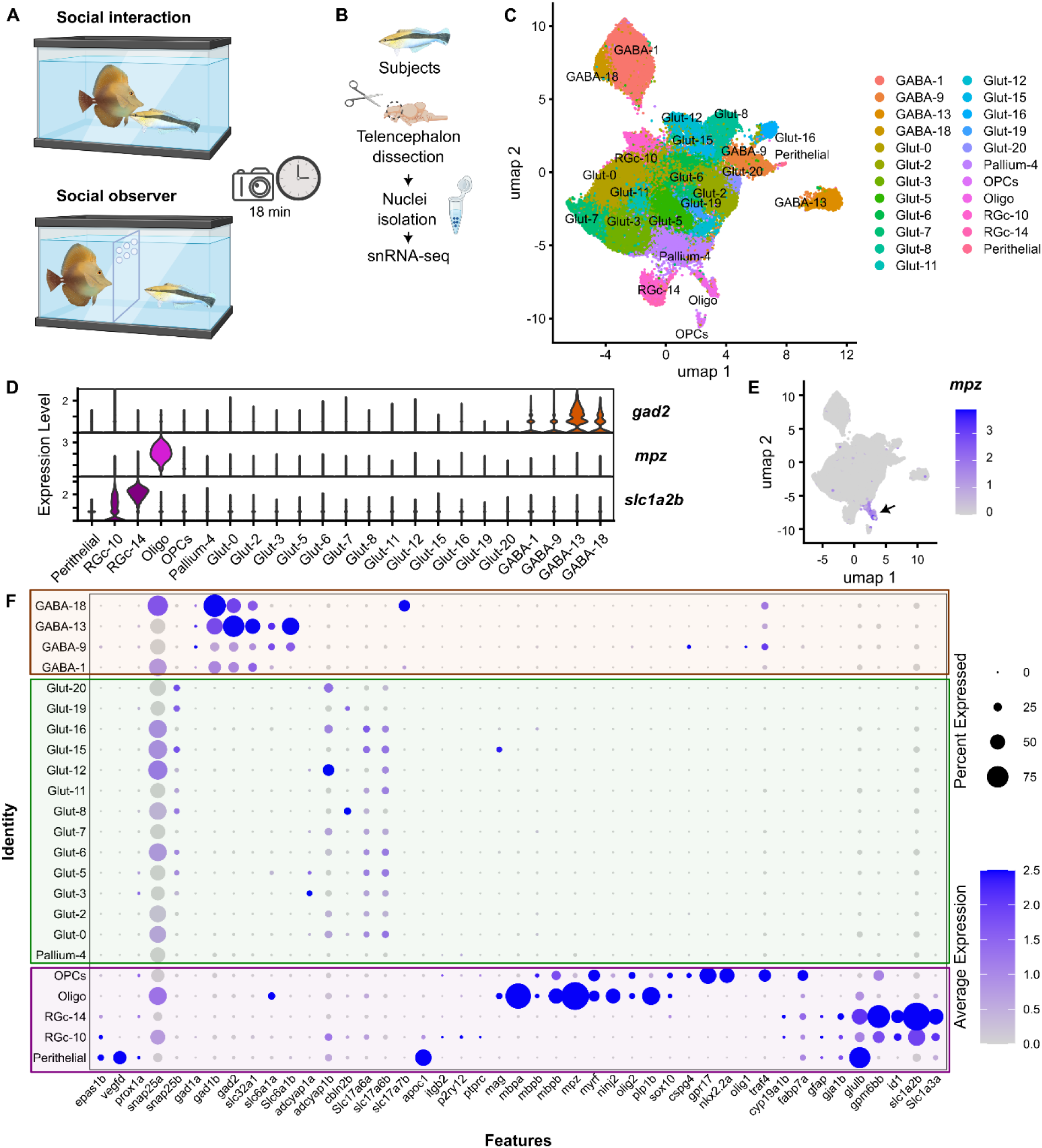
Cell composition of *L. dimidiatus* telencephalon. **(A)** Schematic representation of the behavioural test; *Labroides dimidiatus* (cleaner, on the right side of the tank) and *Zebrasoma scopas* (client, on the left side of the tank) were allowed to interact (social interaction, *n = 12*) or separated by a transparent partition with holes (social observer, *n = 13*) for 18 minutes. **(B)** Simplified representation of dissection protocol for snRNA-seq (*n = 3* subjects per each treatment). **(C)** Integrated UMAP with cell clusters (*n = 23*) assigned. **(D)** Violin plot of selected markers gene expression profiles used for the identification of GABAergic, Oligodendrocytes and Radial Glial clusters. **(E)** Example of the expression profile of the *mpz* marker gene within the UMAP space. **(F)** Dotplot showing the percentage of nuclei and the average expression of genes used to assign cluster identities. GABAergic clusters (orange), Glutamatergic/Pallial clusters (green), non-neuronal clusters (purple). For marker gene reference list see Table S4.

## RESULTS

### Telencephalic nuclei architecture reflects major brain cell classes

Cleaner wrasse telencephalons (*n = 6* individuals, 3 per condition) were dissected, and nuclei were processed and sequenced for snRNAseq, producing 67,207 high-quality nuclei (Fig. 1, B and C; Table S1A and S1B; fig. S1 and S2). Cell composition was assessed and the molecular signature quality of the 23 identified clusters was evaluated by inspecting the percentage of nuclei expressing the top ten marker genes (Table S2; fig. S3). All cell clusters were present in both treatments at similar proportions (fig. S4, A and B). Marker genes associated with the identified clusters were enriched (q < 0.05) for several Gene Ontology categories, including cognition, adult behaviour, regulation of glutamate signalling pathway and dendrite development (Table S3), indicating that these biological processes are the principal features distinguishing cell clusters.

Next, cell clusters were assigned to major neuronal and non-neuronal classes (Fig.1, C to F) using the expression profile of canonical cell type specific marker genes reported in the literature (Table S4). Neurons were broadly classified as GABAergic (*gadb1*^+^, *gad2*^+^, *slc32a1*^+^), or glutamatergic (*slc17a6*^+^, *adcyap1b*^+^). One neural cluster was identified as “Pallial-4” since it had pallial features, but no specific canonical marker genes were expressed (Fig. 1 F; Table S2, S4). Among the non-neuronal populations, we identified clusters expressing canonical markers of oligodendrocytes (*mbp*^+^, *mpz*^+^, *olig2*^+^, *myrf*^+^, Fig. 1E), oligodendrocyte precursor cells (i.e. OPCs; *olig2*^+^, *cspg4*^+^), radial glial cells (i.e. RGc; *gfap*^+^, *cyp19a1b*^+^) and a small population assigned as endothelial-derived cells expressing lymphatic endothelial markers (Perithelial; *vegfd*^+^) (*28*). The RGc clusters, expressed several marker genes commonly associated with mammalian astrocytes (*slc1a2b*^+,^ *fabp7a*^+^, *glula*^+^) (*29*) (Fig. 1D), supporting the hypothesis that radial glial cells in fish share functional similarities with mammalian astrocyte and can therefore be considered astroglia cells (*30*), despite lacking the characteristic stellate morphology.

Finally, as a complementary line of investigation, we performed an unbiased reconstruction of the transcriptional relationship among the identified cell clusters. Cluster tree reconstruction analysis grouped cell types according to their transcriptional similarities. Consistent with their assigned identities, cell populations of the same type clustered together closely with high bootstraps giving support to the cluster subdivision (fig. S5).

### Molecular signatures of social behaviour in cell clusters

To further provide biological insight into the cell clusters and identify transcriptional signatures of behaviourally relevant functions, we generated a curated list of genes specifically associated with neuromodulatory systems and social behaviour in fish, including genes encoding neuropeptides and small molecules neurotransmitter pathways (Fig. 2A; Table S5). From the expression profile, the neuronal cluster GABA-9 showed the most intriguing results. It revealed the co-expression of *nkx2.1*, *lhx6* and the cholinergic markers *lhx8a*, *gbx1* and *isl1* (Fig. 2A). The co-expression of these genes has previously been considered a molecular signature of neurons specifically involved in social behaviour in zebrafish (*Danio rerio*) (*31*). Therefore, GABA-9 likely represents a specialized neural population able to co-synthesizes acetylcholine and GABA associated with social behaviour. Notably, GABA-9 was also characterised by the expression of genes involved in dopaminergic signalling, including dopa decarboxylase (*ddc*), dopamine receptor D2-like (*drd2l*) and tyrosine hydroxylase (*th*) along with genes involved into the serotonergic signalling pathway, such as tryptophan hydroxylase 1 (*tph1a*) and the serotonin receptor (*htr4*), as well as the oxytocin (*oxt*) and oxytocin receptor (*oxtra*). These neuromodulators are known to govern the motivation to interact in cleaners (*18*, *19*), giving further support to the involvement of these cells in the social response. Another interesting feature of this cluster was the expression of *sst1.1* along with *npy*, a gene encoding for the neuropeptide Y. This neuropeptide has previously been associated with feeding behaviour in cleaner fish, where it drives the motivation to find clients and initiate cleaning interactions in wild (*32*). The combined expression of *npy* and *sst1.1* further suggest that this cluster may represents somatostatin^+^, neuropeptide Y^+^ interneurons. Together, these findings highlight the multifunctional neuromodulatory profile of GABA-9 cluster in support of its role as a neural cell population specialized in the regulation of social behaviour.

**Fig. 2.**
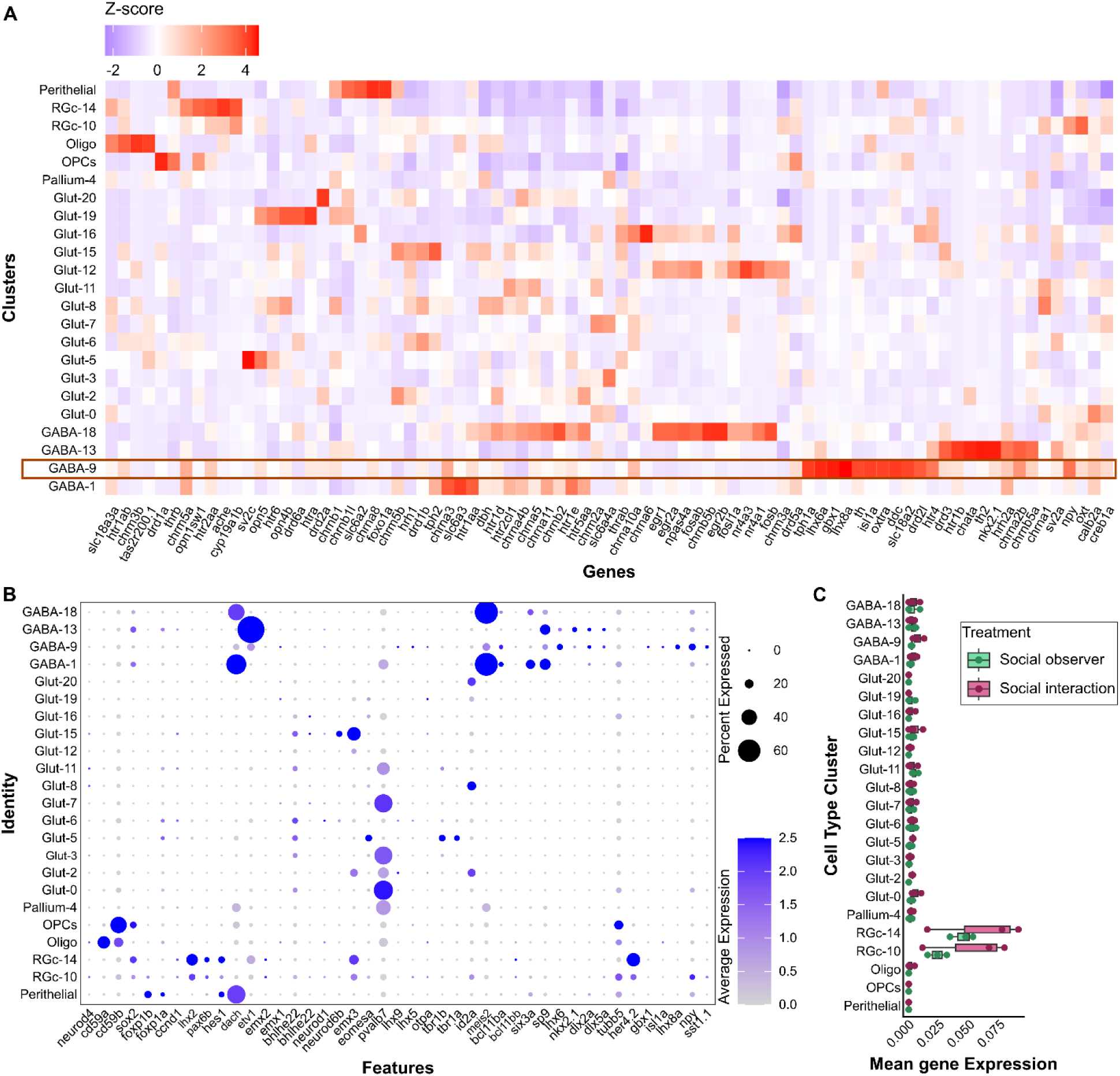
Molecular signatures of cell clusters. **(A**) Heatmap of *z*-score expression profile of selected genes involved in social behaviour in the *L. dimidiatus* snRNA-seq dataset. The orange box highlights the GABA-9 cluster. For the marker gene reference list see Table S5. **(B)** Dotplot showing the percentage of nuclei and the average expression profile of established region-specific neuroanatomical marker genes across all identified clusters. For the gene marker reference list see Table S6A. **(C)** Mean expression of aromatase (*cyp19a1b*) across all clusters and treatments.

Several additional neuronal clusters expressed genes associated with sensory perception including pathways related to visual sensory stimuli. For example, Glut-19 expressed opsin genes (e.g. *opn4b, opn5*) as well as genes encoding serotonin and dopamine receptors (*htra, htr6, drd6a*). The expression profile of Glut-3 was highlighted by nuclei co-expressing genes involved in serotonin pathway (*htr6*, *htr5aa*, *tph1a, slc6a4*). GABA-18 was also expressed several cholinergic and serotonergic receptors and along with GABA-1, these clusters co-expressed genes involved in dopamine pathway, such as *slc6a3*, together with the nuclear receptor *nr4a1* and *nr4a3*, which are implicated in dopaminergic neurotransmission. Interestingly, GABA-18 and Glut-12 exhibited the strongest and near-exclusive expression of Immediate Early Genes (IEGs), suggesting a highly active or plastic neuronal state within these cells (Fig. 2A). Lastly, the gene encoding for aromatase (*cyp19a1b*), a brain-derived estrogen important for radial glial neurogenesis and highly and exclusively expressed in teleost telencephalon (*29*, *33*), was only expressed in both RGc clusters (Fig.2, A and C), highlighting the putative role of these nuclei in shaping social behaviour neurogenesis.

### Anatomical delineation of clusters is conserved across fish species

To determine the neuroanatomical locations of the identified cell clusters, we examined the expression profile of established neuroanatomically restricted a priori genes of interest (Fig. 2B; Table S6A). This revealed spatial expression patterns consistent with distinct neuroanatomical regions previously characterised in other teleost species (*16*, *34*). Notably, broad regional markers clearly delineated glutamatergic pallial and GABAergic subpallial clusters. Specifically, pallial clusters were defined by the expression of *bhlhe22* and *eomesa*, whereas subpallial clusters predominantly expressed *dlx2a* and *dlx5a* (Fig. 2C; Table S6A). Among the pallial glutamatergic clusters, the strong and predominant expression of the *pvalb7* (Glut-0, Glut-3, Glut-7, Glut-11) is consistent with these cell populations contributing to the dorsolateral pallial area (Dl) (*34*). In addition, Glut-2 and Glut-15 displayed the *eomesa*^−^/*emx3*^+^ expression profile, consistent with the medial zone of the dorsal pallium (Dm), which has been proposed as the teleost homolog of the pallial amygdala (*34*, *35*). Interestingly, the RGc-10 cluster co-expressed canonical markers of neuronal precursors, including *tubb5*, *her4.2*, *emx2*, *emx3*, suggesting its contribution to the formation of pallium cell populations (Fig. 2C) (*34*).

To further investigate the neuroanatomical identity of the cleaner fish cell clusters, we used SAMap, a cross-species mapping approach that accounts for protein divergence even among distantly related species. We compared cleaner fish snRNA-seq cell clusters with previously characterised cichlid snRNA-seq cell types that had been integrated with spatial transcriptomics and cross-species comparison (*16*). Overall, we observed strong correspondence between both neuronal and non-neuronal cell clusters, highlighting broad transcriptional conservation between cleaner and cichlid species (Fig. 3A and B, Table S6B). Among the GABAergic populations, GABA-9 showed the strongest correlation with the cichlid cluster 15.3_GABA. Notably, the corresponding cichlid cluster (15.3_GABA) was previously associated with the mouse medial ganglionic eminence (MGE) and proposed as a derivate of the ventral telencephalic region Vl (*16*) (Fig. 3A). The cleaner GABA-9 cluster also strongly expresses *sst1.1* and *npy* (*16*) further supporting its transcriptional correlation with cichlids and proposed homology to mammalian MGE-derived interneurons. GABA-18 instead mapped specifically to the cichlid cluster 4.2_GABA, whereas GABA-1 showed broad correlation across all the 4_GABA cichlid clusters. These cichlid cell populations are spatially localised to the dorsal nucleus of the ventral telencephalic area (Vd) and the supracommisural nucleus of the ventral telencephalic area (Vs) and expressed genes enriched in mammalian striatal medium spiny neurons (MSNs), including *meis2*, *sp9* and *six3a* (*16*). Consistent with our SAMap results, cleaner fish GABA-18 and GABA-1 clusters also almost exclusively co-expressed *meis2*, *sp9* and *six3a* genes (Fig. 2B), giving support to the hypothesis that these clusters represent teleost homologous of the mammalian striatal MSNs.

**Fig. 3.**
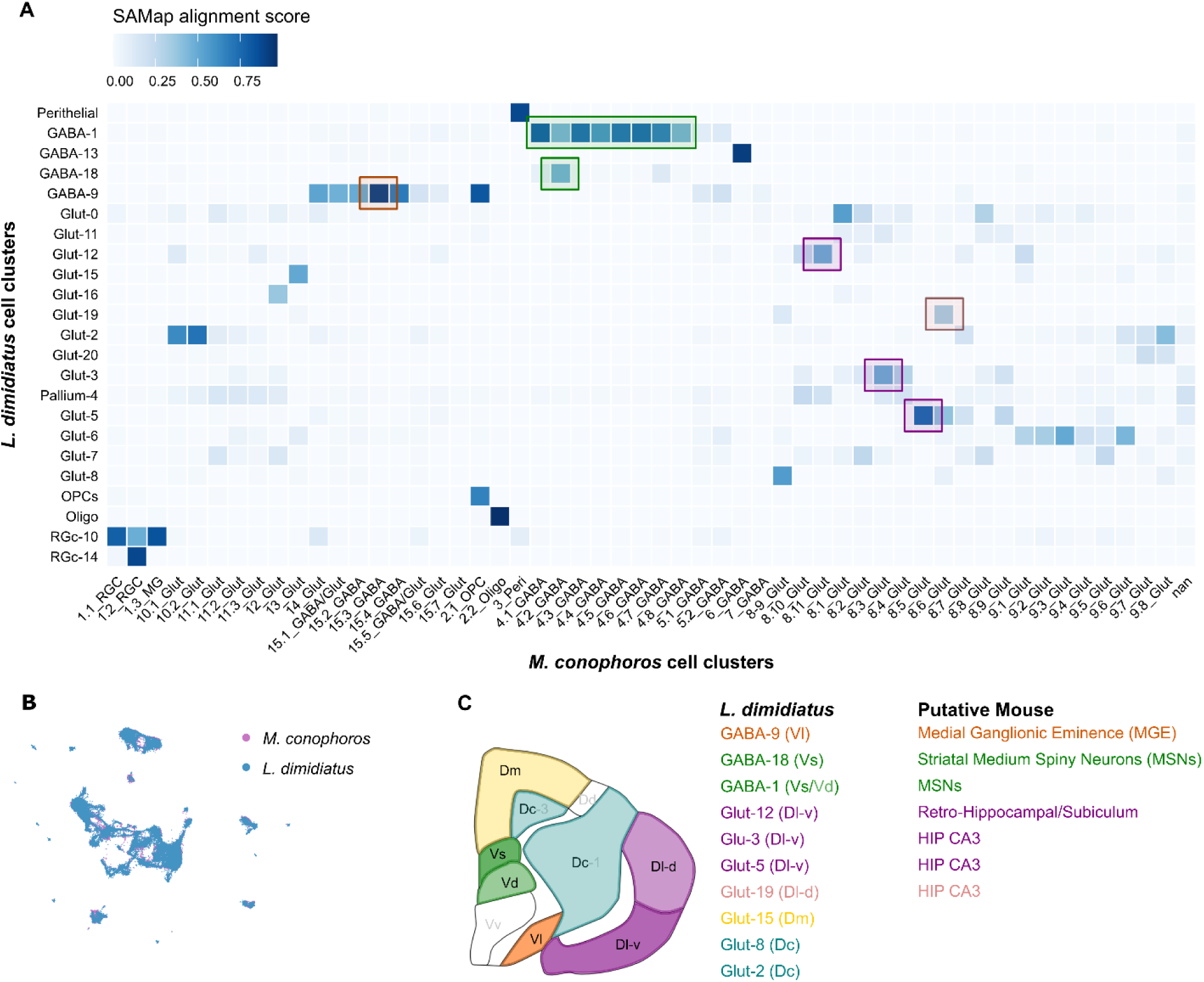
SAMap cell clusters alignment scores between *L. dimidiatus* and *M. conophoros*. **(A**) Mapping of cell clusters between the snRNA-seq *Labroides dimidiatus* (cleaner fish) and *Mchenga conophoros* (cichlid) datasets. Darker colours indicate higher SAMap mapping scores. Coloured boxes indicate putative mouse homologous brain regions. **(B)** SAMap projects of the telencephalic cleaner nuclei (n = 67,207) and cichlid nuclei (n = 33,895) together in UMAP space (cleaner nuclei are ordered to the front). **(C)** Schematic representation of cleaner’ telencephalon coronal section showing assigned tentative regional identity and the corresponding putative mouse brain identity inferred from cichlid dataset (*16*).

Among glutamatergic populations, Glut-19 aligned with the cichlid cluster 8.6_Glut, which was spatially located in the dorsal zone of the lateral subdivision (Dl-d) (*16*). In contrast, Glut-12 showed the strongest correlation with cichlid clusters 8.11_Glut, located within the ventral zone of the lateral telencephalon (Dl-v) (Fig. 3, A and C). Interestingly, this Dl-v associated cichlid clusters corresponded to the mouse subiculum cell type (*16*), which is a structure within the retrohippocampal formation that mediates hippocampal-cortical communication (*36*). Whereases, Glut-3 and Glut-5 aligned strongly with the cichlid cluster 8.3_Glut and 8.5_Glut, respectively. These two cell clusters were putatively located within the Dl-v region and corresponding to the mouse hippocampal CA3 cell-types (*bhlhe22*^+^) (*16*). These correspondences further supports the hypothesis that these teleost cell populations share transcriptional signatures with mammalian hippocampal regions (*34*, *37*). Finally, additional cleaner fish pallial glutamatergic clusters strongly corresponded with specific cichlid cell populations (*16*). Glut-15 aligned closely with cichlid cluster 13_Glut, which spatially maps to the dorsal medial telencephalon (Dm) (Fig. 3, A and C), while Glut-0 and Glut-16 corresponded with cichlid 8.1_Glut and 12_Glut cluster, respectively, spatially localised in the cichlid granule subregion of the dorsal lateral pallium (Dl-g).

### Cleaners do socially interact

Socially interacting cleaners were allowed to interact with the client, while social observers had only visual and olfactory access to the client through a transparent perforated barrier but no direct access (Fig.1A). Socially interacting fish engaged 37 ± 29 times with a client in our behavioural assay (up to 78 times per cleaner), with 12.03 ± 11.6% of time spent interacting (i.e. close body inspection and cleaning activity to remove damaged tissue or scales) and an average duration of 2.93 ± 1.3s. Of those interactions, on average, 3.75 ± 1.7s corresponded to cleaners actively performing cleaning behaviour (i.e. actively removing dead scales or tissues from clients). During interactions, cleaner dishonesty was highlighted by an average of 2 ± 4 of jolts, resulting in cleaners to spend 14.51 ± 15.6% of the interaction providing tactile stimulation for reconciliation, while clients posed an average of 17 ± 12 times per event. In the social observer treatment, the videos were inspected for abnormal or stress behaviour. No such behaviours were observed, and all fish were inspecting the transparent barrier displaying motivation to engage with clients. All behavioural data can be found in Table S7A and S7B.

### Neural excitation is associated with distinct cell clusters

We first investigated transcriptional signatures of neural excitation by analysing the expression of conserved IEGs. Since sn/scRNA-seq approaches generally show low recovery of individual IEG transcripts (*38*), we identified genes selectively co-expressed with each of three well established IEGs (c-*fos*, *egr1* and *npas4*) independently across all clusters (see Method section). This analysis identified a total of 19 IEG-like genes (Table S8), of which 11 were previously known IEGs. We also identified five IEGs that had previously been identified only in cichlids (*39*) and had not been described as canonical IEGs (*DNAJB5, ADGRB1, ITM2C, RTN4RL2*), further supporting the hypothesis that these genes may be specific of teleost species. In addition, we identified three IEGs that have not been previously listed as canonical IEGs but emerged as candidate early genes in our analysis (*MIDN, NAB2, SDC2*). This approach clearly identified two neuronal clusters, GABA-18 and Glu-12, as exhibiting the highest levels of IEG expression (fig. S6). Notably, these cell clusters were putatively assigned to the teleost homologous of the mammalian striatal MSNs and the subiculum, which is implicated in cognitive and behavioural functions (*36*). IEG expression was predominant in both socially interacting and observer fish, with no significant differences among cell clusters (q < 0.05). IEG expression in neurons is associated with neural activity states and synaptic plasticity (*40*). Since cleaner fish from the observer treatment had visual and olfactory access to the client and only active cleaning was excluded, IEG expression is likely driven by the integration of multiple social cues and observer cleaners have an increased neural activation in response to the sensory processing and the attentive brain state.

### Cellular proportion increase in RGc subpopulations in socially interacting fish

Radial glial cells are the principal source of adult neurogenesis teleost (*29*). While the relative cell proportion of cell clusters remained unchanged by social interaction (Table S9A; fig. S7) we hypothesise that behaviourally driven neurogenesis might instead trigger dynamic changes in radial glial cells. To test this, we re-clustered radial glial cells and identified seven subclusters (from RGc-0 to RGc-6; *res = 0.5*; Fig. 4A). Interestingly, the interaction treatment significantly increased the proportion of nuclei present in RG-3 and RGc-4 subclusters (q < 0.05) (Fig. 4B; fig. S8; Table S9B). Furthermore, RGc-3 subcluster showed strong preferential expression of *cyp19a1b*, the gene encoding for aromatase enzyme and specifically involved in radial glial neurogenesis (*29*), with socially interacting fish displaying a propensity to have higher gene expression values compared to social observer fish (fig. S9). Interestingly, the RGc-3 subcluster occupied a transitional region of the UMAP space (Fig. 4A), further supported by the trajectory analysis (Fig. 4B). This subcluster was characterised by the expression of marker genes typically associated with proliferating radial glial cells (e.g. *her4.2*) (fig. S10) (*41*) and therefore potentially representing actively differentiating cells. Taken together, the increased proportion of nuclei in the socially interacting fish along with the *cyp19a1* expression, can partially explain the involvement of radial glial subcluster in the neurogenesis of social behaviour in cleaner fish.

**Fig. 4.**
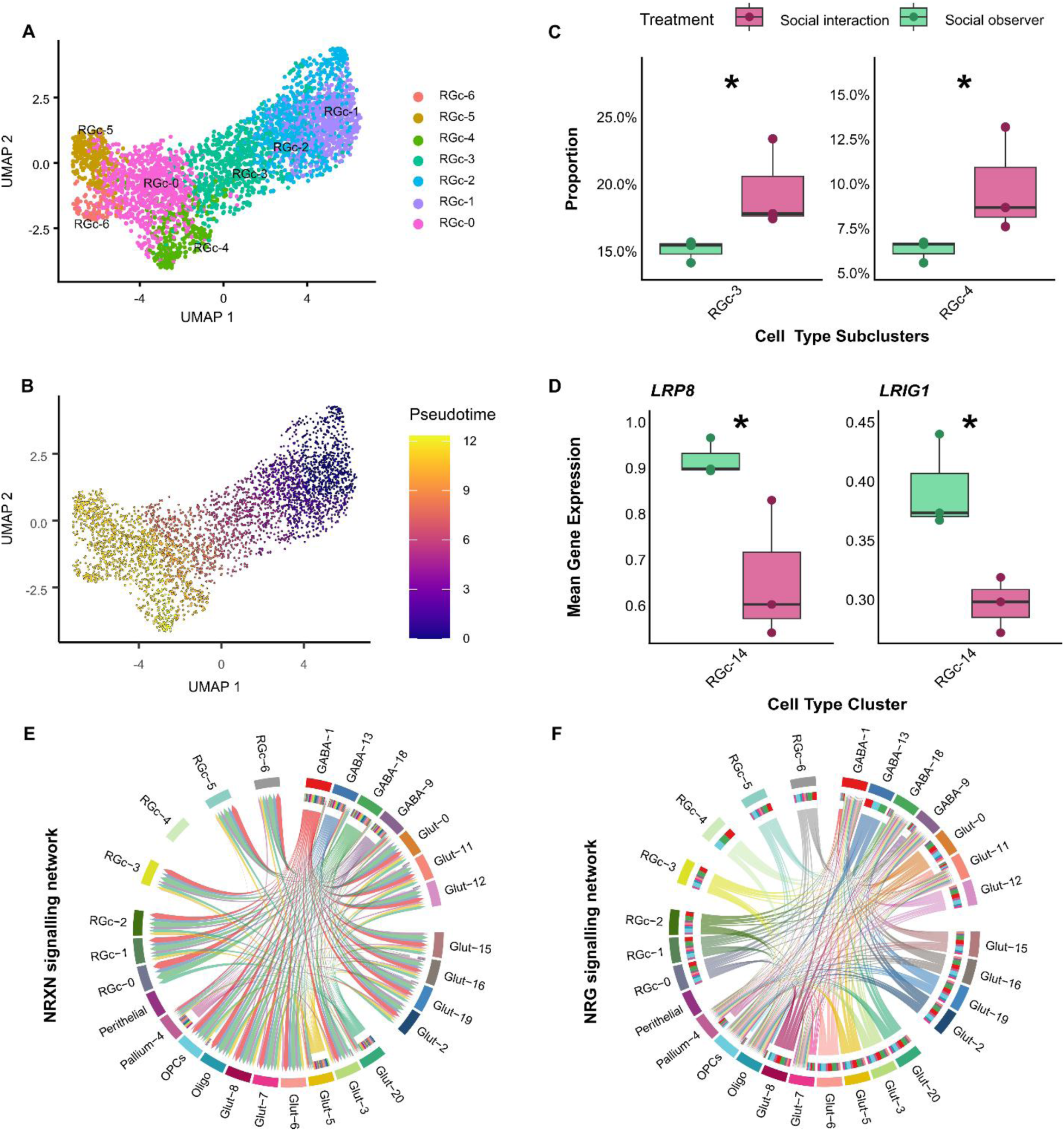
Social interaction elicits different cellular responses. **(A**) UMAP showing the RGc subclusters. **(B)** UMAP showing the RGc subclusters coloured by pseudotime. RGc-1 subcluster was manually assigned as root based on the expression of canonical marker genes (fig S10). **(C)** Cell cluster proportion in RGc subclusters between socially interacting (pink) and social observer (green) treatments. Only significant clusters shown (RGc-3: p = 0.0043, q = 0.025; RGc-4: p = 0.0073, q = 0.025). See Supplementary Material (fig. S8) for the full plot with all RGc subclusters. **(D)** Comparison of DEGs within RGc-14 cluster between socially interacting (pink) and social observer (green) treatments (LRP8, q = 0.024; LRIG1, q = 0.041). Gene expression values were aggregated to pseudobulk by calculating the mean expression per gene for each cell type within individual biological samples. **(E-F)** Chord diagrams illustrating predicted cell-cell communication networks involving **(E)** Neurexin-Neuroligin ligand-receptor interactions and **(F)** Neuregulin-Erbb4 signalling pathway.

### Predicted cell interactions identify directional cell communications

Heterogeneous cell populations communicate through different neural circuits, and neurogenesis is regulated by both neural activity and social experiences (*42*). We hypothesise that the increased proportion of radial glial cells observed in interacting fish could reflect enhanced neural activity. We performed a cell-cell communication analysis using CellChat (*43*) and examined the predicted communication patterns among cell clusters, with particular emphasis on radial glial subclusters and on neuronal clusters for which we had previously inferred putative anatomical locations. Intercellular communication modelling identified GABA-18 and GABA-1 clusters as the signalling hubs within the network (Fig. 4E; Table S10), with GABA-9 and Glut-12 acting as the major target receiver clusters. Among the radial glial subclusters, RGc-3 exhibited the highest level of outgoing signalling (Table S10). Mechanistically, this communication network was driven by Neurexin-Neuroligin ligand-receptor interactions, followed by the Neuregulin-Erbb4 pair (Fig.4, E, F), suggesting an underlying architecture of synaptic connectivity and functional neuronal communication.

### Social interactions elicit transcriptome responses in distinct cell populations

Social interaction in cleaners induced changes in gene expression in specific cell populations. We analysed the Differentially Expressed Genes (DEGs) for each cell cluster by comparing cells from social observers and socially interact fish using MAST. Despite the presence of vision and smell sensory stimuli in the social observer treatment, the analysis revealed 13 DEGs (FDR < 0.05), indicating that these genes are likely specific to the active cleaning interaction (Table S11). Among these genes, the radial glial cluster RGc-14 revealed the downregulation of the leucine-rich repeats and immunoglobulin-like domains protein 1 (*LRIG1*), apolipoprotein E receptor 2 (*ApoER2*, also called low-density lipoprotein receptor-related protein 8, *LRP8*) (Fig. 4D) and lipin 2 (*LPIN2*) in socially interacting fish. Interestingly, *LRIG1* is a well-known stem cell marker that regulates stem cell quiescence (*44*) whereas *LRP8* is involved in the signal transduction of the reelin (*RELN*), an extracellular glycoprotein important for brain synaptogenesis, dendrite maturation, synaptic plasticity and neurotransmitter release (*45*, *46*).

Among the neuronal clusters, for example, GABA-1 showed the downregulation of the calcium channel, voltage-dependent, R type, alpha 1E subunit critical for neurotransmitter release, synaptic plasticity and cellular excitability as a potential response to the interaction event, along with the upregulation of the gamma-aminobutyric acid (GABA) A receptor, beta 2 (*GABRB2*), and the protein tyrosine phosphatase, receptor type, U, a (*PTPRU*), which is predicted to be involved in signal transduction and neuron projection development (Table S11).

### Co-expression networks correlate with social behaviour and anticipatory state

Given the complexity of snRNA-seq data, we used the high-dimensional weighted gene co-expression network analysis (hdWGCNA), a framework designed to infer co-expression networks from single cell datasets (*47*), to identify coordinated gene expression networks associated with social behaviour. Based on the identity of cell clusters and the expression of social behaviour related genes, we focused our analysis on GABA-9 and Glut-12 neuronal clusters individually, and GABA-1/GABA-18 combined. To characterise the biological relevance of the identified co-expression modules, we constructed the cluster dendrogram (Fig. 5A), computed Module Eigengenes (MEs) at the single cell level and estimated eigengene-based connectivity (kME) for each gene. By looking at the GABA-9 cluster, the identified modules included several hub genes (Table S12) with known roles in neuronal communication and plasticity. For example, the GABA-9-M5 module included *BAI3*, a gene involved in axonal and dendritic arborizations as well as excitatory synapse formation (*48*, *49*), and the neuroligin-1 (*NLGN1*), a key regulator of excitatory synapses and long-term memory (*50*). Similarly, the GABA-9-M1 module revealed hub genes associated with glutamatergic signalling, such as *GRIK3*, as well as *NPY*, and the synaptotagmin Ia (*SYT1*), a protein involved in synaptic vesicle release from photoreceptors (*51*). Furthermore, the differential expression analysis identified GABA-9-M5 and GABA-9-M1 significantly positively and negatively correlated, respectively (Fig. 5B), in socially interacting fish compared to observer cleaners, emphasizing the distinct transcriptional programs within the GABA-9 neuronal cluster elicited by the social interaction (Table S13A).

**Fig. 5.**
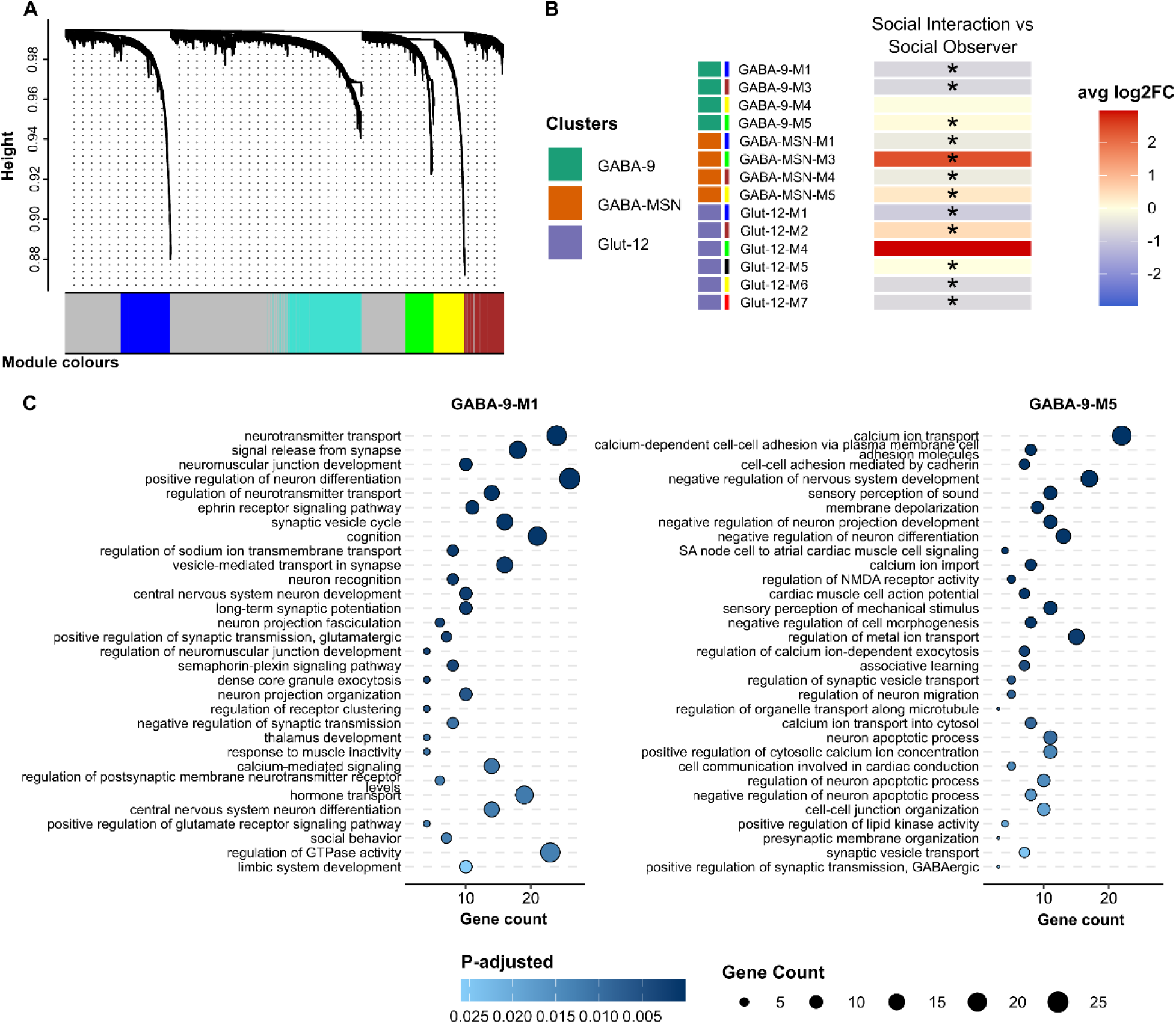
Gene co-expression modules architecture. **(A)** Example of hdWGCNA dendrogram showing GABA-9 cluster. **(B)** Module eigengene (ME) heatmap showing the average expression of GABA-9 (green), GABA-MSN (orange), and Glut-12 (purple) co-expression modules correlated to the social behaviour treatment. Higher values (red) represent positive correlation with the social interaction treatment. Significant values (q < 0.05) from the “differential module eigengene” analysis are reported with “*”. The corresponding module colours assigned by hdWGCNA are reported on the side of the module names. **(C)** Top 32 exclusive enriched biological processes within GABA-9 modules (M1 on the left side and M5 on the right side) ranked by adjusted p-value (q < 0.05). Non-redundant GO terms were determined using “REVIGO” (*53*) with default parameters.

GABA-9-M1 and GABA-9-M5 shared several enriched biological processes related to synaptic transmission, signalling and adult behaviour (e.g. GO:0035249, GO:0048167, GO:0030534) (Table S14). Despite the similarities, among other terms, GABA-9-M1, which was positively associated with the observer cleaners, was exclusively enriched in cognition and memory (GO:0050890, GO:0007613), limbic system development and long-term synaptic potentiation (GO:0021761, GO:0060291) along with positive regulation of neuron differentiation and hormone transport (GO:0045666, GO:0009914), highlighting a module with genes strongly connected to a preparedness and attentive state of cleaners (Fig. 5C; Table S14). On the other hand, GABA-9-M5, which was positively correlated with cleaners actively engaging in social interactions, displayed exclusive enrichment in biological processes related to associative learning (GO:0008306), calcium ion transport and synaptic vesicle transport (GO:0006816, GO:0048489) along with the regulation of NMDA receptor activity (GO:2000310) (Fig. 5C; Table S14). Notably, molecular functions in GABA-9-M5 were predominantly associated with terms centred around channel activity (including GO: 0099094, GO:0099106, GO:0016247), calcium signalling and glutamate receptor activity (e.g. GO:0099604, GO:0098988), emphasizing the active transmission machinery existing in GABA-9-M5 module (Table S14). Finally, integration of co-expression network analysis with GSEA further supported the biological relevance of the identified modules, revealing a significant (padj < 0.05) broad overrepresentation of metabolic processes and gated channel activity in GABA-9-M5 (Table S15A). Together, these findings suggest the existence of distinct gene regulatory networks within GABA-9 neuronal cluster in response to social behaviour stimuli.

Glut-12 cluster was putatively assigned to the subiculum mammalian cell types (Fig. 3A and C), suggesting a potential role in the regulation of a broad range of behavioural functions. Consistent with its presumed function, the Glut-12 modules were enriched for genes associated with locomotory behaviour and learning or memory functions (e.g. GO:0007626, GO:0007611) (Table S16 and S17). Notably, the Glut-12-M2 module, which was positively associated with the socially interacting fish (Table S13B), was exclusively enriched for functions related to presynapse organization, presynaptic active zone organization, and the regulation of adenylate cyclase-activating G protein-coupled receptor signalling pathway (GO:0099172, GO:1990709, GO:0106070) (Table S17). These enrichments were absent from any of the GABAergic modules investigated, highlighting the specificity of the Glut-12-M2 module in processes involved in presynaptic assembly and involved signalling pathways.

Lastly, both GABA-1 and GABA-18 (named as “GABA-MSN”) were putatively assigned to the MSNs mammalian cell types (Fig. 3A and C), an area expressing dopaminergic receptors and involved into emotional states behaviours (*52*). Consistent with its presumed function, these GABAergic clusters were enriched for genes associated with learning or memory (GO:0007611) (Table S19 and S20). Notably, the GABA-MSN-M1 module, which was positively associated with the social observer fish (Table S13C), was exclusively enriched for functions related to olfactory bulb interneuron differentiation (GO:0021889) and mesenchyme development and differentiation (GO:0060485, GO:0048762) while GABA-MSN-M5 module, which was positively associated with the socially interacting fish (Table S13C), displayed an enrichment in molecular functions related to dopamine receptor binding (GO:0031749, GO:0050780) (Table S20). These enrichments were absent from any of the GABA-9 or Glut-12 modules, delineating the specificity of these modules in regulating different aspects of social behaviour in cleaner fish.

## DISCUSSION

Here we provide the first cellular atlas of the telencephalon of a model for advanced social cognitive abilities, the cleaner wrasse *Labroides dimidiatus*. Characterising telencephalic cell types enabled a detailed examination of the cellular organization of the telencephalon in a highly socially interacting fish, uncovering strong conservation of cell populations across teleost species and identifying candidate cellular components and molecular pathways involved in the regulation of social behaviour.

Among the identified cell types, our study revealed a cholinergic-GABAergic neural cell cluster (GABA-9) with a transcriptional profile suggestive of a role in social behaviour. This cluster expresses *nkx2.1*, *lhx6*, and the cholinergic markers *lhx8a*, *gbx1* and *isl1*, a transcriptional signature closely matching a recently discovered zebrafish (*Danio rerio*) telencephalic neuronal population required for social orienting behaviour and defined by the co-expression of GABAergic and cholinergic markers (*31*). This molecular signature is also characteristic of Medial Ganglionic Eminence (MGE)-derived neuronal cells in the developing mammalian forebrain, where the coordinated activity of *lhx6*, *lhx8a*, *isl1*, and *nkx2.1* is essential for MGE specification and neuronal development (*54*, *55*). In vertebrates, *lhx6* and *lhx8* are key regulators of GABAergic and cholinergic cell fate, respectively, and are predominantly expressed in the ventral telencephalon, particularly the MGE, which gives rise to several interneuron classes, including somatostatin-expressing interneurons (*54–56*). The cleaner fish’ GABA-9 transcriptional profile is highly consistent with these mammalian cell populations. In particular, the co-expression of *sst1.1* and *npy* suggests that these cells represent a putative MGE-derived somatostatin^+^/neuropeptide Y^+^ interneuron class (*34*). This interpretation is further supported by the comparison of single cell and spatial transcriptomic data from cichlids, indicating localisation within the ventrolateral (Vl) telencephalon (*16*). Together, these findings support the hypothesis that GABA-9 cells are homologous to MGE-derived neurons and represent an evolutionarily conserved subpallial neuronal population. Notably, GABA-9 cells also express genes associated with dopaminergic and serotonergic signalling pathways, two neuromodulatory systems known to regulate social behaviour and learning in cleaner fish (*19*, *26*). Furthermore, *npy* has recently been linked to feeding behaviour and reward-related dopaminergic pathways in cleaners (*32*). The coexistence of cholinergic, GABAergic, dopaminergic and serotonergic signalling components within the same neuronal cluster suggests that these cells may function as an integrative hub coordinating multiple neuromodulatory pathways involved in social behaviour. Collectively, our results identify a conserved neuronal cell cluster that may play a key role in the modulation of cleaner fish social behaviour and represents a promising target for future functional studies.

Our neuroanatomical characterisation is broadly consistent with findings from cichlids (*16*, *39*), zebrafish (*34*) and goldfish (*Carassius auratus*) (*37*). We identified strong transcriptional homologies between cleaner fish neuronal cells and cell types located in homologous subpallial and pallial brain regions across teleosts, despite potential variation arising from differences in experimental protocols and cell type annotations. Among the GABAergic cell types, the cleaner GABA-18 and GABA-1 clusters exhibited transcriptional and anatomical similarity to cichlid Striatal Medium Spiny Neurons (MSN)-like cell types spatially localised to the ventral telencephalon (*16*), a region considered homologous to the mammalian nucleus accumbens within the striatum (*17*) with similar correspondences have been reported in zebrafish (*34*) and goldfish (*37*). The dorsal nucleus of the ventral telencephalic area (Vd) is a brain region that receives substantial dopaminergic input and is considered homologous to the mammalian nucleus accumbens of the striatum (*17*). Given the central role of mammalian MSNs in dopamine-dependent behavioural regulation (*57*, *58*), these GABAergic cell clusters may similarly contribute to the regulation of social behaviour in cleaners. In agreement with these findings, GABA-MSN modules showed broad enrichment in functions related to learning or memory and exclusive enrichment in molecular functions related to dopamine receptor binding relative to socially interacting fish, strengthening the presumed functional role attributed to these cell types and their involvement in social behaviour.

Among glutamatergic clusters, the cleaners Glut-12 was most transcriptionally similar to cichlid cell types located within the ventral subdivision of the dorsolateral pallium (Dl-v), a region proposed to be homologous to the mammalian retrohippocampal formation, including the subiculum (*16*). Comparable relationship have also been reported in zebrafish (*34*) and goldfish (*37*). The hippocampal formation, including subiculum, is highly conserved across vertebrates and plays a central role in a wide range of behavioural functions, including spatial navigation, learning and memory as well as synaptic plasticity and long-term potentiation, thereby promoting memory formation and retrieval (*36*, *59*, *60*). Consistent with these results, behavioural studies have implicated fish Dl-v in hippocampal-like functions, including learning and memory (*61*). In agreement with these findings, Glut-12 modules displayed broad enrichment in functions related to locomotory behaviour, learning or memory and specific enrichments within module 2 (Glut-12-M2) in presynapse organization and presynaptic active zone organization positively correlated to socially interacting fish. Furthermore, Glut-12 exhibited a strong immediate early gene expression profile, indicative of elevated neuronal activity (*40*), suggesting a role in information processing and memory formation during social interactions. Together, these findings suggest that the transcriptional and anatomical similarities between cleaner fish and other teleost neuronal cells may reflect conserved functions related to social behaviour, specifically in learning and memory. Nevertheless, the extent to which the cleaner fish Dl-v area is homologous to the mammalian hippocampal formation remains to be fully elucidated. Further investigation of Dl-v cellular composition, connectivity and functional properties will be necessary to clarify its role in learning and memory formation in cleaner fish.

Social experiences, including interactions, learning, and memory, can influence adult neurogenesis, the process by which new neurons are produced from neural stem cells and integrated into existing brain circuits (*42*). Consistent with this, we observed an increased cell proportion within two radial glial subclusters (RGc-3 and RGc-4) in socially interacting fish. This aligns with the extensive capacity of teleost adult neurogenesis compared with mammals (*62*). In cichlids, for example, radial glial cells have been proposed to mediate neuronal production and circuit remodelling in response to complex social behaviours (*39*). Several lines of evidence further support a neurogenic role for these cells in interacting cleaner fish. The aromatase-expressing glial subpopulations identified may contribute to the regulation of both cell proliferation and social behavioural responses, as disruption of aromatase signalling alters telencephalic proliferation in zebrafish (*63*). In addition, inferred directional cell-cell communication involving the neuregulin-Erbb4 signalling pathway identified, suggests active regulation of neuroblast proliferation, migration, and differentiation by radial glial, consistent with established roles of neuregulin signalling in mammalian neurogenic niches and the subventricular zone (*64*, *65*). Further support comes from the RGc-14 cell cluster, which exhibited downregulated expression in response to social interaction in *LRIG1*, a negative regulator of neurogenesis in the mammalian ventricular-subventricular zone (*44*). Given that loss of *Lrig1* enhances neurogenic activity in mice (*44*), the pattern identified may indicate increased neurogenesis in radial glial cells following active social interaction. We also observed downregulation of *LRP8*, a key receptor in the Reelin signalling pathway (*66*) that mediates adult neuroplasticity and synaptogenesis (*45*, *46*, *66*). It is tempting to speculate that this change may reflect compensatory feedback associated with pathway activation during social interaction. Although direct experimental evidence linking social behaviour, radial glial activity and neurogenesis in cleaner fish require further exploration, our results suggest that social interactions induce molecular changes consistent with enhanced neurogenic processes, highlighting this species as a promising model to investigate the mechanisms underlying socially induced neurogenesis in vertebrates.

Previous work with bulk-RNA showed that social interactions induce changes in forebrain gene expression in cleaner fish (*67*), raising the possibility of identifying specific cell clusters involved in mutualistic social behaviour. By isolating direct social information from observer cleaners, we identified transcriptional changes not only within radial glial cells (*LRIG1*, *LRP8*), but also within neuronal clusters. GABA-1, for example, showed differential expression of calcium channel subunits such as the voltage-dependent, R type, alpha 1E subunit. Calcium channels, together with glutamatergic signalling, are key regulators of learning, memory processes and synaptic plasticity in cleaners and are considered major molecular drivers in cleaning interactions (*20*, *32*, *68*). Interacting cleaners also exhibit upregulation of *PTPRU*, a gene codifying for a receptor-type tyrosine phosphatase, involved in signal transduction, neuronal projection development and neuronal maturation (*69*). This GABA-1 cluster was identified as one of the major signalling sources through the neurexin-neuroligin pathway, known to play a central role in synaptic transmission and development (*70*), which further strengthens the putative involvement of this GABA-1 cell cluster as important mediator of social information processing in socially interacting cleaners.

Beyond the signalling capabilities of other identified neuronal cell clusters, the GABA-9 cluster emerged as a promising integrative hub coordinating multiple neuromodulatory pathways during social interactions. Co-expression gene network module 5 (GABA-9-M5) was positively correlated with socially interacting fish and enriched in hub genes involved in neural network formation and memory processes, including *BAI3* and *NLGN1*. Variants in *BAI3* have been linked to neuropsychiatric disorders such as schizophrenia and bipolar disorder (*48*), highlighting its importance in higher cognitive functions. *NLGN1*, in turn, is a key regulator of synapse formation, synaptic development and plasticity, and memory retention through mechanisms involving N-methyl-D-aspartate (NMDA) receptor signalling (*50*). Consistent with this, regulation of NMDA receptor activity was uniquely enriched within this module, suggesting that active social interaction may be mediated through this pathway. Given that glutamatergic signalling is a major molecular driver of learning, memory, and cleaning interactions in cleaner fish (*20*, *32*, *68*), it is intriguing to speculate that *NLGN1*-mediated modulation of NMDA receptor activity can support the synaptic currents required to establish and maintain prolonged interactions with clients. Reinforcing its role in social responses, GABA-9-M5 was enriched for calcium signalling, glutamate receptor activity, and gated channels. Together, these glutamatergic and calcium - dependent pathways likely drive enhanced synaptic transmission, cognitive flexibility, and social behaviour in cleaner fish (*20*). Notably, the exclusive enrichment of associative learning processes within this co-expression network module is particularly relevant in the context of cleaner-client interactions. Cleaners preferentially feed on client mucus, a dishonest behaviour that often elicits client jolts followed by chasing (*14*, *71*). However, cleaners also have remarkable learning and discriminative abilities, and they can resolve this conflict by adjusting their cleaning service (*72*). In our experiment, chasing events consistently followed client jolts, providing opportunities for cleaners to learn from negative feedback and adjust their service quality (*73*). Together, these findings suggest that cells within the GABA-9-M5 module may contribute to the establishment and maintenance of social interactions, as well as in behavioural adjustments that promote the formation of future social memories.

While GABA-9-M5 emerged as a key module for socially interacting fish, GABA-9-M1 module was positively correlated with social observers. Within this module, key hub genes, such as synaptotagmin Ia (*SYT1*), suggest the role of visual information in shaping the transcriptional responses of observer cleaners exposed to indirect social stimuli through synaptic vesicle release from photoreceptors (*51*). From this analysis, another mechanism emerged as central regulator in social observers. *NPY* is one of the most abundant orexigenic neuropeptides in the vertebrate brain (*74*) and has previously been linked to cleaner motivation to seek clients and initiate cleaning interactions in the wild (*32*). Through its modulation of reward-related pathways, including the mesolimbic dopaminergic system (*75*), *NPY* is thought to influence motivational states associated with food-seeking behaviour (*32*). Given that this cleaner fish species relies on client’s ectoparasites as primary food source (*10*), the association of *NPY* with observer fish further supports its role in promoting the motivation to engage in cleaning interactions. More broadly, these findings point to a central role for the GABA-9-M1 module in regulating motivational and reward-related processes. Reinforcing this interpretation, GABA-9-M1 was uniquely enriched for cognition, long-term synaptic potentiation, neuron differentiation, and limbic system development. Because limbic circuits govern emotion, motivation, and reward (*75*) these pathways suggest an attentive, preparatory state preceding social interactions. Furthermore, this mesolimbic integration, combined with *NPY* as a major hub gene, likely drives an adaptive anticipatory state in observer cleaners that enhances exploration and attention to potential clients. Overall, GABA-9-M1 primes the neural circuitry for subsequent cleaning behaviour through integrated sensory, motivational, and cognitive processing.

Several limitations of this study should be acknowledged. First, although snRNA-seq provides unprecedent transcriptomic resolution, it lacks the spatial information. To mitigate this limitation, we integrated our cellular atlas with cross-species neuroanatomical comparison. However, future studies using spatial transcriptomics will be necessary to resolve the spatial organization and dynamics of neuronal and glial cell populations. Second, while we identified associations between differential gene expression, co-expression modules, and cell types associated with specific social stimuli, causal relationships remain to be established. Lastly, while it is intriguing to speculate that cell proliferation and neurogenesis contribute to social behaviour in cleaners, additional experimental evidence is needed to determine the extent and functional significance of these processes within the complex social behavioural context of this species.

Collectively, this study suggests that GABA-9 neuronal cells may represent a key cellular substrate underlying the molecular mechanisms of complex social behaviour in *L. dimidiatus*. Here we show how socially active cleaners exhibited distinct transcriptional and gene co-expression network signatures compared with social observers, reflecting experience-dependent behavioural adjustments at the cellular level. Radial glial cells also emerged as important contributors to social behavioural responses. By integrating snRNA-seq, cross-species neuroanatomical comparison, differential gene expression and gene co-expression analysis, we identified a reproducible molecular signature linking specific cell types to social behaviour. These findings on cleaner fish sociality provide a single cell perspective on the neural cells and molecular mechanisms underlying social behaviour and establish a framework for investigating the neural basis that govern complex social interactions across vertebrates.

## MATERIALS AND METHODS

### Fish husbandry

Adult cleaner wrasses *Labroides dimidiatus* (i.e. cleaner fish) and the client surgeon fish *Zebrasoma scopas* (*76*), were obtained from a local fish store (Sealife Hong Kong Limited) and transported to The Swire Institute of Marine Science (SWIMS) aquarium. Similarly to other studies on the same species (*11*), fish were acclimated for 18 days to the aquarium conditions before any behavioural experiment with conditions similar to their natural environment (salinity at 35 ppt, temperature 25°C, pH 8.1) and fed ad libitum once per day with mysis frozen shrimps (Hikari Bio-Pure®) for cleaners and spirulina (Tropical®) for clients. Each cleaner fish was kept in a separate individual tank (30 L) to avoid aggression among individuals, whereas one to three clients were kept together in each 50 L aquarium. Individual tanks had a semi-closed flow-through aquatic system where alkalinity levels, dissolved carbons and pH were daily monitored and kept constant. Tanks were kept under a photoperiod of 12h/12h (light/dark cycle) and temperature was automatically controlled. Ammonia, nitrite and nitrate levels were checked twice a week using colorimetric tests (Sera test). Additional handheld equipment was used to complement the automatic systems with a manual daily monitoring of seawater temperature, dissolved oxygen (Pro 2030), salinity (portable refractometer), and pH (pH meter, BWA-2018SD). Individual observational tanks were used for testing the behaviour and water parameters were manually checked before the start of any experiment. A summary of water chemistry and tanks parameters can be found in the Supplementary Material Table S21A.

### Behavioural experiment

Behavioural tests were performed in independent observation tanks (50 x 30 cm and 25 cm high; 50 L) in an isolated room. On the day of the experiment, cleaner wrasse fish were placed into an observational tank with either (1) cleaner-client pair (social interaction) or (2) non-interacting fish (social observer) (Fig. 1A). Fish were first acclimatized to the observation tank for 20 minutes prior to the experiment by adding an opaque partition. Then, the opaque partition was raised, and behaviour was recorded by a video camera (GoPro Hero 10 black) positioned on the front of the tank. In the social interaction treatment cleaner and client were allowed to interact, while in the social observer treatment, cleaners had visual and olfactory access to the client through a transparent perforated barrier, allowing us to isolate the effects of indirect social information in the absence of active cleaning interactions. Similarly to previous interaction test conducted on cleaner fish (*77*), each trial was recorded for 18 minutes to have a good representation of the behaviour and the clients never engaged with other cleaners more than twice per day. Clients used for each experimental treatment were similar in size and colour pattern (according to the fish supplier catalogue were 6-10 cm size) and never interacted with more than one cleaner fish per day. Cleaners underwent a 24 h fasting period before the interaction to avoid gene expression patterns related to food metabolism (*67*).

Behavioural analysis for the interaction treatment were measured as reported in previous studies (*20*, *67*, *78*) investigating both motivation to interact and interaction quality of cleaners. Briefly, motivation was measured by counting the number and proportion of interactions initiated by cleaners (i.e. close body inspection and cleaning activity to remove damaged tissue or scales) and the ratio of clients “posing” display conducted to attract the cleaner (i.e. client posing display/time not interaction). Whereas interaction quality was evaluated based on the duration of interaction events, the proportion of interaction time spent in performing tactile stimulation (i.e. cleaners touching with pectoral fins the client to reduce stress and enhance interaction duration), and the frequency of client jolts per 100s of interaction (signals of cheating/dishonesty by the cleaners). For observer fish, behaviour was inspected for the presence of any abnormal behaviour such as erratic movements or aggressive postures and active interest towards the transparent barrier (i.e. close inspection of the barrier).

All videos were analysed using the event logging software “Boris” v8.27.10 (*79*) with the behavioural catalogue detailed in Supplementary Table S7B. Both cleaner and client fish were considered as a focal subject in the analyses. A total of 25 cleaners were included for the behavioural analysis (*n = 13* social observers; *n = 12* social interaction).

### Tissue sampling and nuclei isolation

Following the behavioural experiment, cleaner wrasses were immediately euthanized by severing spine (*n = 3* social observers*; n = 3* social interaction), measured for standard length and body weight, and brain were extracted from the skull (Table S21B). All procedures were carried out in approval of the Committee on the Use of Live Animals in Teaching and Research (CULATR) of the University of Hong Kong and by the Department of Health of the Hong Kong Government (Protocol and Ref. numbers: #22-051, #24-676), following the CULATR guidelines and regulations as well as the ARRIVE guidelines. Telencephalons, excluding olfactory bulbs, were dissected under a stereo microscope in ice cold Neurobasal medium (with 2% B27 and 0.5 mM Glutamax; Thermo Fisher Scientific) containing 0.2 U/μl RNase Inhibitor (Sigma). Dissected telencephalons were rapidly frozen in liquid nitrogen and stored at -80°C.

Nuclei were isolated following a protocol adapted from the 10x Genomics Demonstrated Protocol CG000393 • Rev A (https://cdn.10xgenomics.com/image/upload/v1660261285/support-documents/CG000393_Demonstrated_Protocol_Adult_Mouse_Nuclei_Isolation_RevA.pdf) and optimized for *L. dimidiatus* telencephalon. Briefly, 1.5 ml of lysis buffer containing 10 mM UltraPure Tris-HCl, pH 7.5 (Invitrogen), 10 mM NaCl (Invitrogen), 3 mM MgCl_2_ (Invitrogen), 0.1% NP-40 Surfact-Amps™ Detergent Solution (Thermo Scientific), 1% BSA (Sigma), 0.2 u/ul of Protector RNAse inhibitor (Roche) and RNase-free water were added to each frozen telencephalon tissue (*n* = 3 per behavioural condition) and incubated on ice for 2 minutes. Following the incubation, tissue was mechanically triturated using a pestle and the lysate was fully dissociated using a P1000 pipette and left on ice for other 10 minutes. Dissociated tissue was transferred to a 40 μm filter (Genever®-CS04) to remove large debris and centrifuged (600 rcf; 5 min; 4°C). The supernatant was discarded, and the pellet was resuspended in 1.5 ml chilled wash and resuspension buffer containing dPBS (Gibco) with 2% BSA (Sigma), 0.2 u/ul of Protector RNAse inhibitor (Roche). Nuclei suspension was then filtered through a 10 μm filter (pluriSelect) to remove small debris and nuclei aggregates and centrifuged (500 rcf; 5 min; 4°C). Purified nuclei were resuspended in 150 μl of wash and resuspension buffer to reach a final concentration between 700-1900 nuclei/μl, counted in a disposable haemocyte (Millicell®) using the Ethidium Homodimer-1 (Thermo) staining dye at the fluorescent microscope (Nikon DS-Ri2 camera) and then immediately processed to the 10x Genomics platform.

### Single-nuclei RNA sequencing

The isolated nuclei suspensions were processed with 10x Genomics Chromium Next GEM Single Cell 3ʹ Reagent Kits v3.1 (Dual Index; User guide GC000315, Rev F) and Chromium Next GEM Chip G Single Cell Kit with a target recovery of 10,000 nuclei per sample. Single nuclei were then encapsulated into Gel-Beads-in-emulsion (GEM) by 10x Chromium iX. Library were constructed and sequenced on an Illumina NovaSeq 6000 (paired ends, 150 bp) with a total throughput of ∼100Gb at the Centre for PanorOmic Science (CPSO), the University of Hong Kong.

FASTQ files were processed with 10x Genomics Cell Ranger v6.0.1 pipeline (*80*), and reads were aligned to the *L. dimidiatus* genome assembly (ASM3071049v1) using a splice aware alignment tool (STAR) within Cell Ranger and by including introns (*--include-introns*). Gene annotations were custom-made obtained from the same assembly (see Supplementary text “Genome Annotation Pipeline” for a detailed description). The final *Labroides dimidiatus* annotation file used in this study can be retrieved at “https://www.schunterlab.com/resources”. Cell Ranger filtered out UMIs that were homopolymers, contained N, or contained any base with a quality score less than 10. Overall, we sequenced a total of 76,199 nuclei at an average depth of 30,406 reads/nuclei. On average, reads mapped confidently to both genome and transcriptome were 85.1% and 38.6%, respectively (Table S1A).

### snRNAseq data pre-processing and normalization

Data was first quality inspected and pre-processed for each library independently in R v4.5.2. Ambient RNA reads were first removed using SoupX v1.6.2 (*81*) with default parameters. Low quality cells were then filtered (nFeatures above 250 and below 5000; nCount < 18000; % of mitochondrial genes < 3), and doublets were removed using scDblFinder v1.24.10 (*82*) using default settings. Quality control metrics for each library are provided in the Supplementary Material (Table S1A and S1B). In total, 67,207 cells passed the quality filters and were processed for downstream analyses.

Data were imported into Seurat v5.4.0 (*83*) for data normalization according to the SCTransform v2 pipeline with a Gamma-Poisson Generalized Linear Model and by regressing out the percentage of mitochondrial reads (*84*). Linear dimensional reduction was independently conducted for each sample using the “RunPCA” function with 50 principal components (PCs). After Elbow plot inspection, we included 30 PCs for the Uniform Manifold Approximation and Projection (UMAP) by using the “RunUMAP” and “FindNeighbours” functions.

### Data integration, clustering and cell marker identification

Samples were integrated using the “IntegrateLayers” function from Seurat v5.4.0 using the reciprocal PCA analysis (RPCA). Clusters were computed using “FindCluster” function at different resolution employing the Clustree v0.5.1 package (*85*) to show the relationship between clusters, and the final selected cluster resolution was 0.8. The biological identities of each cluster were investigated incorporating unbiased analysis of cluster specific marker genes as well as a supervised examination of previously established cell marker genes. Cluster specific marker genes were identified using the “FindAllMarkers” function (*only.pos = TRUE, min.pct = 0.25, logfc.threshold = 0.25, test.use = “wilcox”*) and were then intersected with well stablished cell type specific markers identified from the literature (Table S4) (*16*, *29*, *34*, *37*, *39*, *86*) as well as markers from CellMarkers 2.0 (*87*). To further confirm the transcriptional relationship of the different clusters, we performed a cell type tree reconstruction based on pairwise expression distances between clusters (all expressed genes used) using 10,000 bootstraps (*88*). For the tree reconstruction, a gene was considered expressed only if the average expression was greater than three transcripts per million in at least one of the identified clusters.

### Functional enrichment analysis of cluster specific markers

Functional enrichment analysis of Gene Ontology (GO) categories among cluster-specific marker genes was investigated to gain further insights of the major processes characterising each cluster. The AnnotationForge v3.22 (*89*) package was used to build the genome-wide annotation (OrgDb) containing all specific GO for *L. dimidiatus*. Then, we computed profiles of each gene cluster using the “enrichGO” function provided by ClusterProfiler v4.18.4 (*90*) and FDR-adjustment (*q* < 0.05) were considered statistically significant.

### Ortholog gene assignment

We used diamond v2.1.24 (*91*) to perform BLAST analysis between *L. dimidiatus* sequence and protein sequence of zebrafish (*D. rerio*; genome assembly used: GCF_049306965.1) and mouse (*M. musculus*; genome assembly: GCF_000001635.27) with the following parameters *-f 6, -- sensitive*. In case of one-to-many or may-to-one matches, only the gene with the highest percent identity match was retained. Ultimately this approach returned a set of 18,516 and 16,482 genes for zebrafish and mouse, respectively. For consistency and easier interpretation, we included the zebrafish gene symbol annotation within the manuscript and in all included figures.

### Identification of IEG-like genes and IEG expression analysis

To improve the detection of signatures associated with neural excitation, we adopted a method previously described by Johnson et al., (2023) and Parker et al., (2024). Briefly, we identified genes that were selectively co-expressed with each of three canonical Immediate Early Genes (IEGs), c-*fos, egr1, npas4*, across all clusters and, within each cluster, nuclei were divided into IEG-positive and IEG-negative groups. The function “FindMarkers” from Seurat was then used to assess the differential gene expression between these groups (*logfc.threshold = 0*, *min.pct = 1/ min(cluster_sizes)*, reflecting the size of the smallest cluster). All genes failing to meet these criteria within a cluster were excluded and assigned a p-value of 1. Since FindMarkers requires at least three nuclei in each group comparison, clusters containing fewer than three IEG-positive nuclei were removed from the analysis. Genes detected in most clusters and significantly upregulated (p < 0.05) in IEG-positive nuclei across most of those clusters were considered significantly co-expressed with each canonical IEG. Finally, only genes significantly co-expressed with all three canonical IEGs were classified as IEG-like genes.

To analyse the difference in neural excitation, treatment-associated IEG expression was analysed in each cluster. We assigned to each nucleus an IEG score equal to the number of unique IEG-like genes previously found (n = 19). Treatment differences in IEG score were analysed using a generalized linear mixed effect regression model assuming a negative bimodal distribution (*glmer.nb*) with the “lmer” package in R (*86*). Treatment was considered categorical fixed effect and the individual from which each nucleus derived was treated as random effect (significance with q < 0.05).

### Neuroanatomical inference and cross-species comparison

To investigate the neuroanatomical origins of the cleaner cell clusters, we first interrogate the expression profile of known neuroanatomically restricted a priori genes of interest (e.g. pallial and subpallial marker genes; see Table S6A for a complete list). Secondly, snRNA-seq data from the cichlid *Mchenga conophoros* (*39*) was integrated to allow a comparative analysis of the transcriptional similarities of cell types. We specifically selected this species thanks to the availability of both single nuclei and spatial transcriptomic data, which can therefore be used as a teleost reference for deeper comparative neuroscience investigations. To perform this cross-species comparison, we used SAMap (*92*), a python tool specifically designed for comparative analysis of distantly related species. To determine sequence similarities, we initially downloaded the protein sequence from the same reference genome assembly used to create the snRNA-seq dataset (GCF_000238955.4). Next, we used the SAMap’s map_genes.sh bash script to identify reciprocal blast hits between cleaner and cichlids. Then, we converted into h5ad file format the raw gene counts matrix of both cleaner and cichlids snRNA-seq datasets and finally the data were processed using the run function from SAMap with default settings and by specifying the gene symbols of each protein. To further identify the genes driving cell type relationships between cleaner and cichlids, we used the GenePairFinder function from SAMap with default settings. The function specifically finds genes driving the relationship by using a pairwise comparison of cell types. For our study, we retrieved all the genes found driving the relationship, which are considered to positively contributing to the cross-species correlation of cell types (Table S22).

We considered as homologous all the cross-species mutual top hit cell type pairs. We inferred the neuroanatomical locations based on the present SAMap results as well as the spatial mapping of the cichlid cell populations and the cross-species comparison reported in a previously published study (*16*).

### Cluster proportion analysis

Cluster proportions were compared between social observer and social interaction treatments using a mixed effect binomial regression model employing the “glmer” package in R. Inside the model, treatment was considered a fixed effect, individuals were considered random and the outcome variable was the nuclei that either belonged to the cluster of interest or not. We adjusted p-values with the Benjamini-Hochberg correction Differences and in cluster proportions were considered significant with q < 0.05.

### Re-clustering of Radial Glial clusters

To investigate the radial glial subpopulation diversity, we pooled the RGc clusters (RG-10 and RG-14), and we re-clustered the cell population using the SCTransform workflow with a shallow resolution of 0.5 for cluster identification. We then analysed the subcluster proportion differences between treatments using a mixed effect binomial regression model employing the “glmer” package in R as described above. To test the role of radial glial in cleaner fish neurogenesis, we further analysed the aromatase (*cyp19a1*) expression among the identified subclusters. We used the “glmer.nb” function from lme4 R package. Inside the model, treatment was considered a fixed effect, individuals were considered random and the natural logarithm of library sizes per nucleus was set as an offset to normalise the sequencing depth across nuclei. As above, we adjusted p-values with the Benjamini-Hochberg correction Differences (significant level with q < 0.05).

### Trajectory analysis of RGc-subclusters

Monocle3 (*93*) was used with the recommended default settings to build single-cell trajectories. The root of the pseudotime analysis was manually identified by plotting and exploring the marker genes previously established (Table S4) and expressed within each RGc subclusters. Cells expressing typical markers of undifferentiated cells should represent earlier stages in their developmental trajectory and were set as the root.

### Cell-cell communication analysis

To assess the probability of ligand-receptor interactions associated with specific signalling pathways and directional communication strength (i.e. connection weight) between cell populations, we used the R package CellChat v2.2.0 (*43*). We first identified the mouse orthologs of the *L. dimidiatus* (see “ortholog gene assignment” section) and then created a CellChat object with the “createCellChat” function. The “cellChatDB.mouse” was used to infer cel-cell communication. Over-expressed genes and interactions were identified using the “identifyOverExpressedGenes” and “identifyOverexpressedInteractions”, respectively. The communication probability and cellular network were computed using the default function parameters.

### Differential expression analysis

To identify genes correlated to the interaction treatment, we performed a differential expression analysis with MAST’s two-part hurdle model v1.36 (*94*). Raw UMI counts from the RNA assay were supplied for model construction (mixed effect modelled as follow: *zlm(formula = ∼ngeneson + mt_reads + Treatment + (1 | orig.ident)*) with a similar approach previously reported in other studies (*95*). Cellular detection rate, percentage of mitochondrial reads and treatment level were fitted to the model as covariates, with sample origin fitted as a random effect to overcome pseudo replication bias similar to other single cell studies (*96*). To avoid biases, we removed lowly expressed genes expressed < 10% of cells. Differential gene expression was then computed between social observer and social interaction treatments using a likelihood ratio test and by adjusting p-values with the Benjamini-Hochberg correction, and genes were considered differentially expressed with FDR < 0.05.

### High-dimensional weighted correlation network analysis (hdWGCNA)

High dimensional weighted gene co-expression network analysis (*47*) was performed to identify key genes associated with cooperative behaviour in specific cluster (GABA-9, Glut-12 and the combination of GABA-1/GABA-18, named as “GABA-MSN”). We first selected the soft threshold value by using the “TestSoftPowers” function (GABA-9, softPower of 8; Glut-12, softPower of 9; GABA-MSN, softPower of 6), then the co-expression network was constructed with the “ConstructNetwork” function. Module eigengenes and module connectivity were computed with default parameters, and the first top 20 highly connected genes (hub genes) were sorted by kME and extracted. One of the identified modules for each of the analysed clusters, namely GABA-9-M2, Glut-12-M3 and GABA-MSN-M2, were primarily enriched for genes associated with mitochondria activity and cellular metabolism. As our focus was on molecular pathways related to behaviour and neural signalling, these modules were excluded from further analysis. The identified modules were further investigated for significant positive and/or negative correlations with the treatment by analysing the “differential module eigengene” with the “FindDMEs” function (*min.pct = 0.25*). The biological context of the co-expressed modules was analysed by performing an enrichment test using the “enrichGO” function provided by ClusterProfiler v4.18.4 (*90*) and FDR-adjustment (significance *q* < 0.05). Enriched functions were then inspected using “REVIGO” (*53*) with default parameters to reduce functional redundancies.

### Gene set enrichment analysis (GSEA)

To provide further biological context to the interaction behaviour, we performed GSEA with the fgsea R package v1.37.4 on specific co-expression modules identified by the hdWGCNA analysis (namely, GABA-9-M1 and GABA-9-M5). We supplied a ranked gene list for the specific module of interest, ordered by module eigengene-based connectivity (kME), and used all the remaining genes identified in the co-expression analysis, excluding the grey module, as a background set. GO pathways were retrieved from the *L. dimidiatus* annotation file deposited in “https://www.schunterlab.com/resources”. Functions were considered significative with padj < 0.05.

## Acknowledgments

We would like to thank all people who gave us suggestions and inputs to make this experiment possible, including all the people from the Earlham Institute who never hesitate to help. We also warmly thank Hugo Lee for providing the illustrations of the fish in Figure 1A and Hang Cheong Chan for the precious help with the aquarium facility at SWIMS. We are also grateful to all the members of the Schunter’s Lab for their constant support.

## Author contributions

Conceptualization: C.S., D.D.; Methodology: D.D., C.S.; Investigation: D.D., C.S.; Visualization: D.D. Supervision: C.S.; Writing—original draft: D.D., C.S.; Writing—review & editing: D.D., C.S., D.J.W., W.H, I.M., A.L.

## Data and Material availability

All data needed to ensure reproducibility and evaluate the conclusions in the paper are present in the paper and/or the Supplementary Materials. The raw snRNA-seq data generated for this work are deposited on NCBI Gene Expression Omnibus (accession number GSE347338). Custom codes used are also available from GitHub (https://github.com/Deb96-lab/snRNA_brain_profile).

