## Supplementary Material for "Single-nucleus profiling of social behaviour in a mutualistic reef fish"

**Authors:** Debora Desantis *et al.*

**This PDF file includes:**

Figs. S1 to S10
Supplementary Text
References for supplementary text (1 to 13)

**Other Supplementary Materials for this manuscript include the following:**

Files from S1 to S22

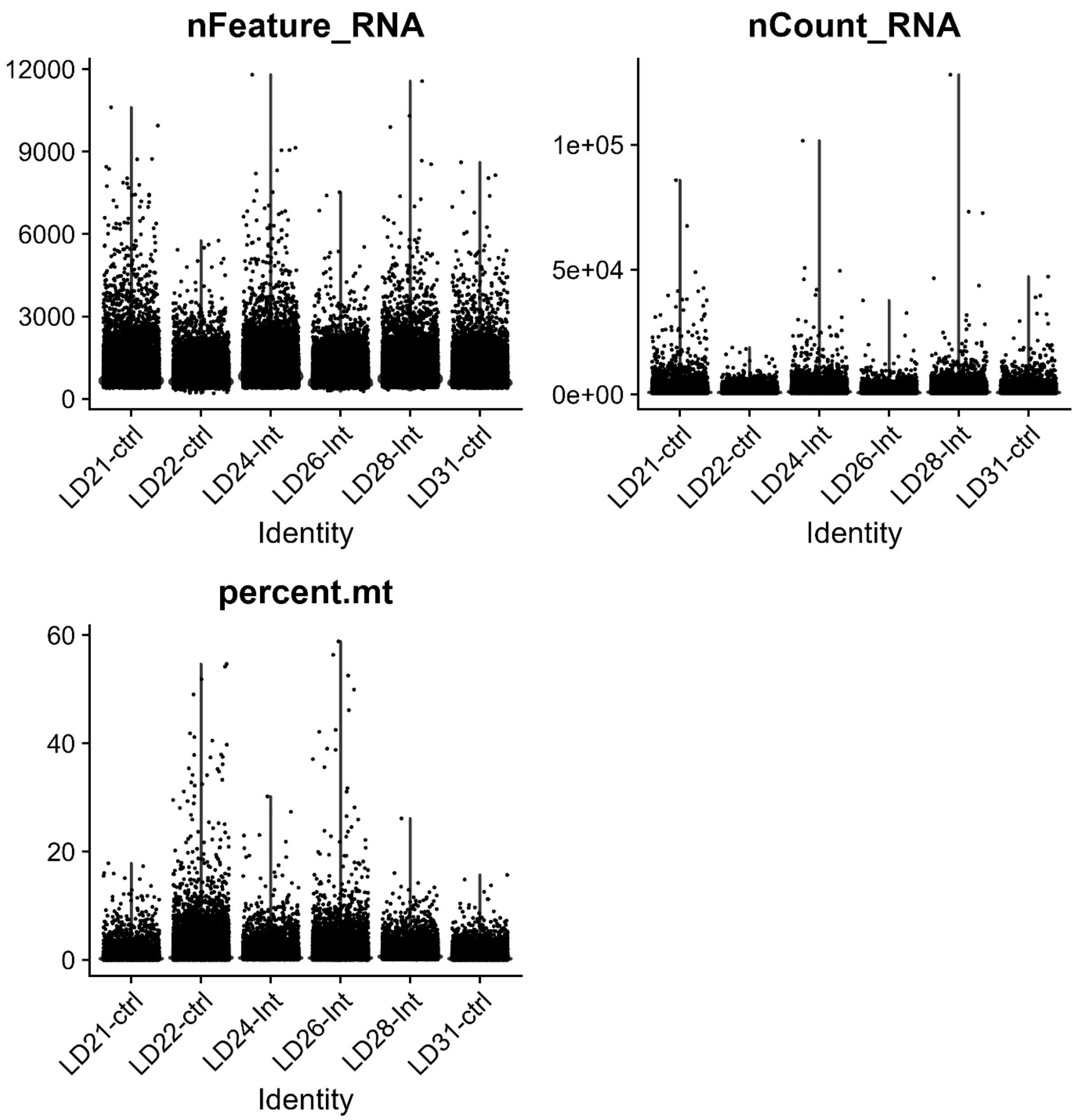


**Fig. S1. Quality control for snRNA-seq data before quality filtering.**

Violin plots showing: left, the number of detected unique genes ("nFeature_RNA"); right, the number of unique molecular identifiers (UMIs, "nCount_RNA"); and bottom, percentage of mitochondrial genes ("percent.mt") in each cell from each sample before data preprocessing. Samples LD24-Int, LD26-Int, and LD28-Int are the "social interaction", while LD21-ctrl, LD22-ctrl, and LD31-ctrl are the "social observer".

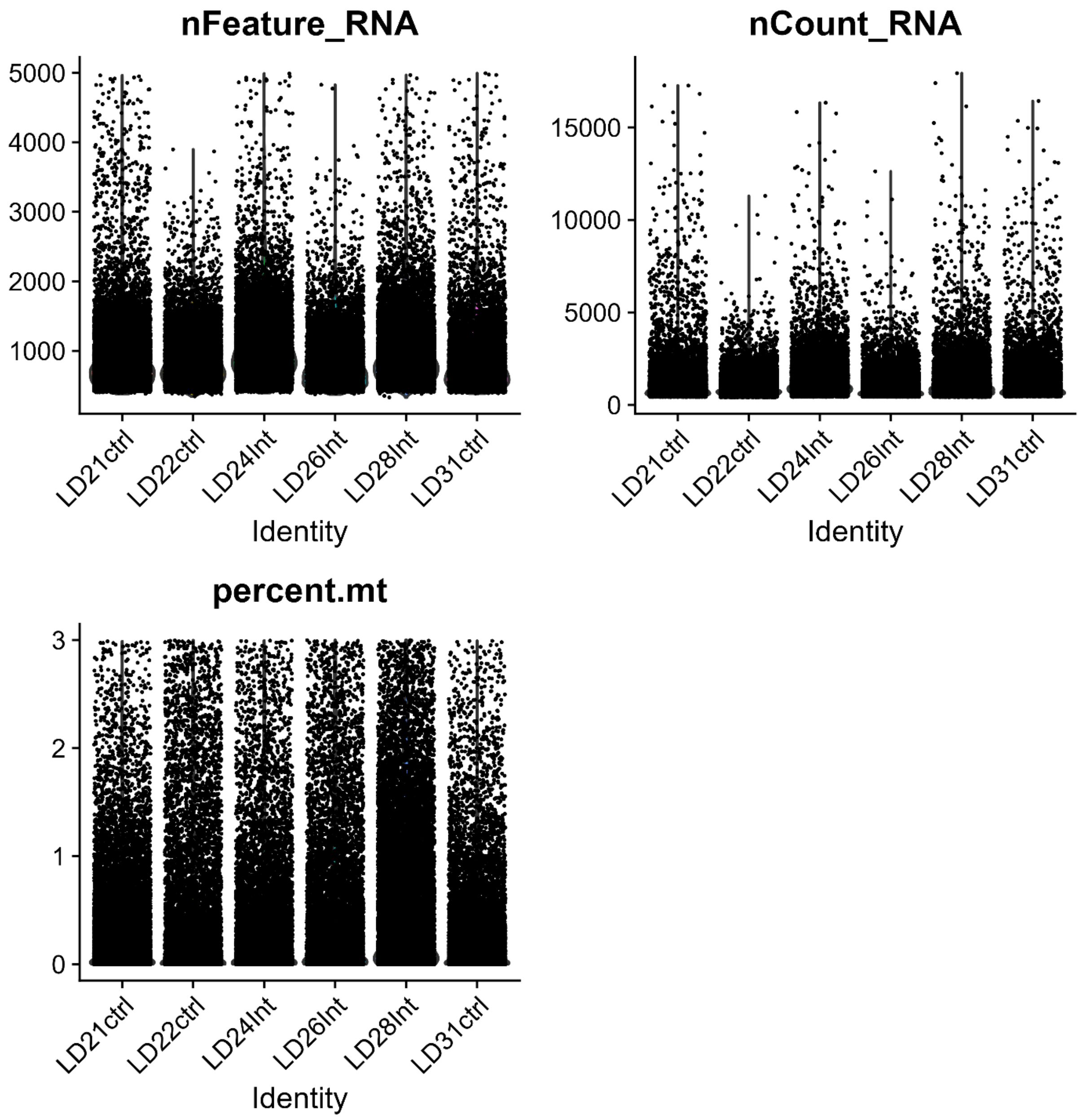


**Fig. S2. Quality control for snRNA-seq data after data preprocessing steps.**

Violin plots showing: left, the number of detected unique genes (“nFeature_RNA”); right, the number of unique molecular identifiers (UMIs, “nCount_RNA”); and bottom, percentage of mitochondrial genes (“percent.mt”) in each cell from each sample after quality control data preprocessing. Samples LD24-Int, LD26-Int, and LD28-Int are the “social interaction”, while LD21-ctrl, LD22-ctrl, and LD31-ctrl are the “social observer”.

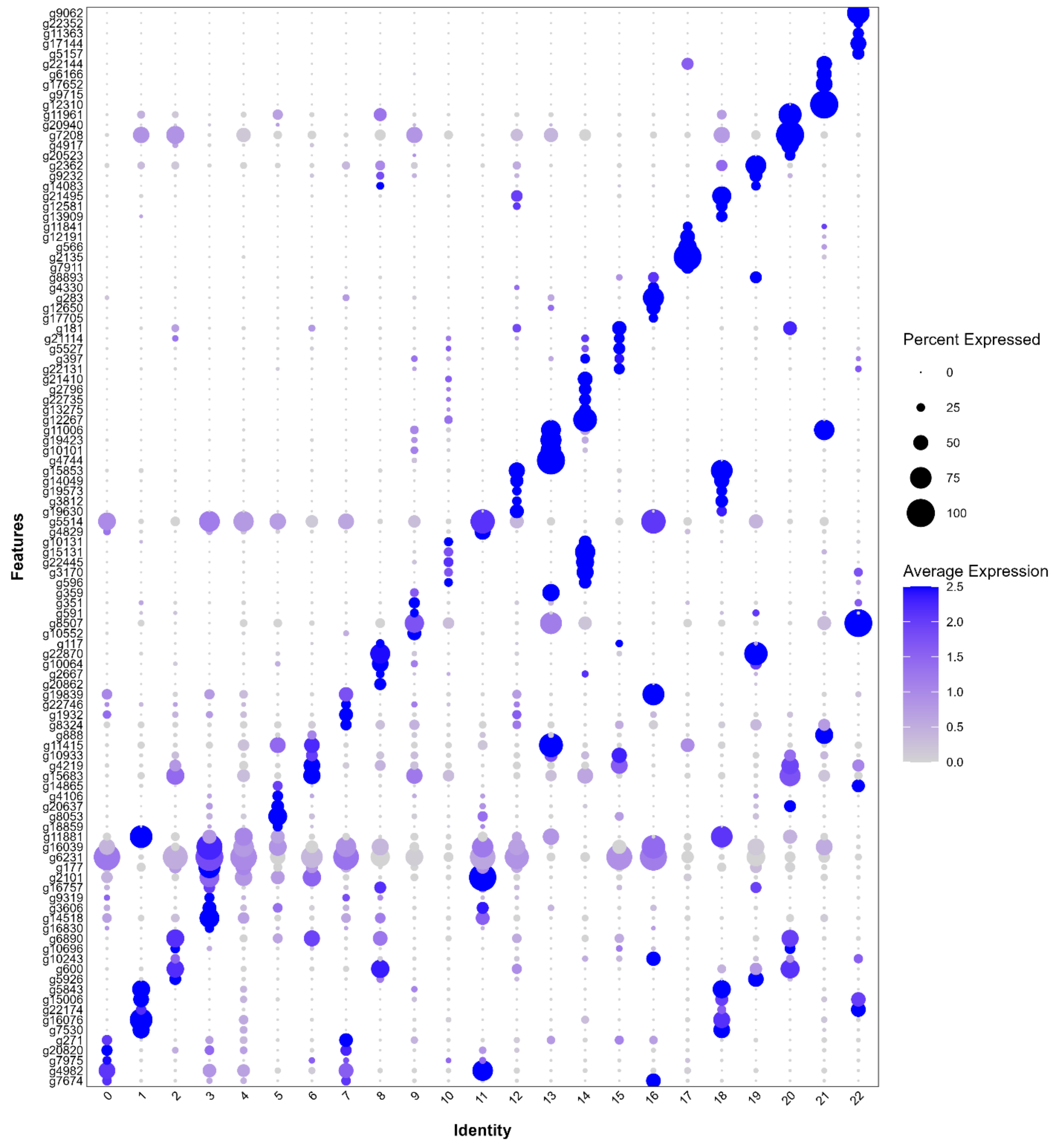


**Fig. S3. Cluster specific marker genes.**

Dotplot representing the percentage of nuclei and the average expression of the top5 marker genes per cluster and cell type family identified from FindAllMarker function. For the corresponding gene labels, see Table S2.

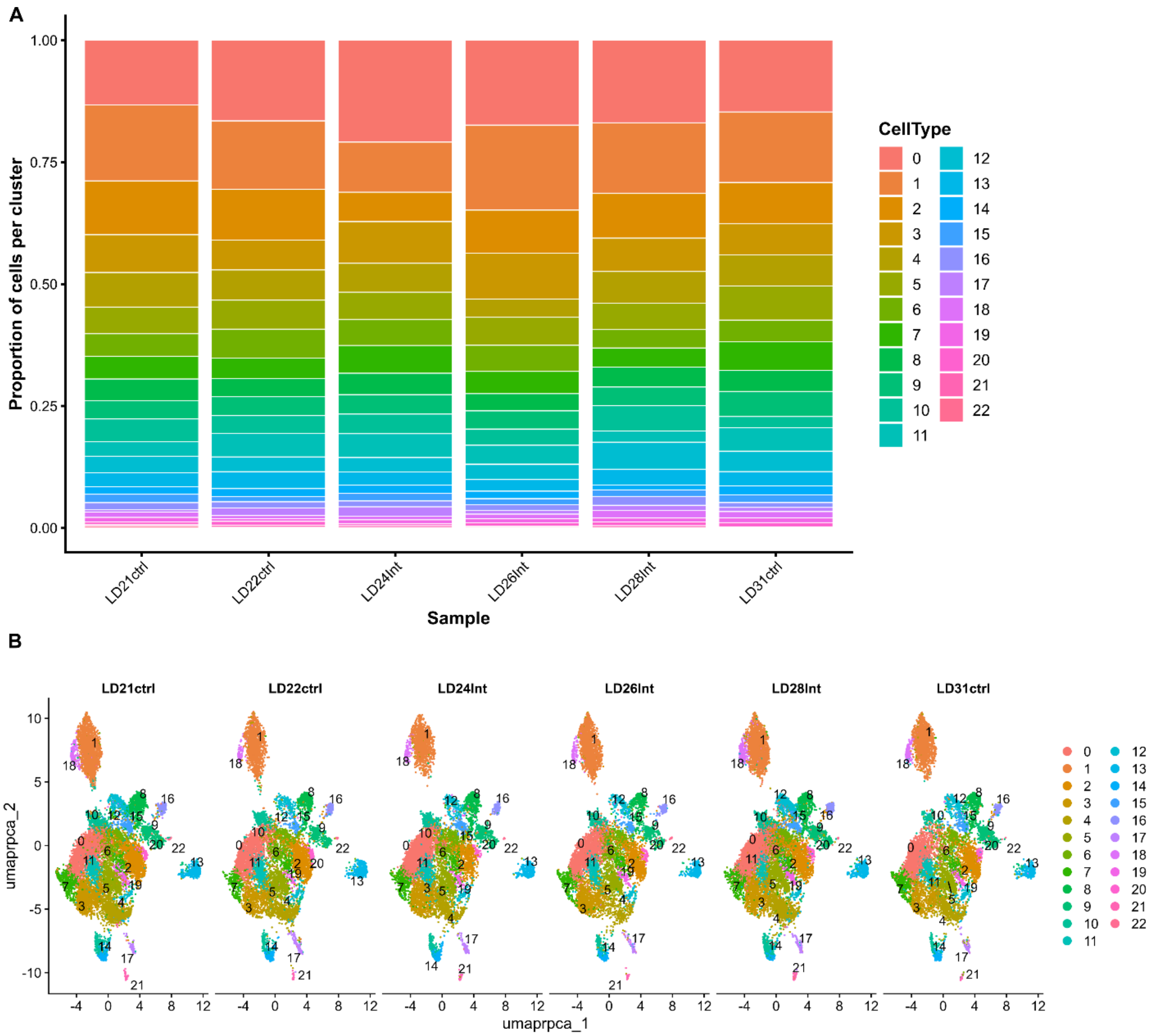


**Fig. S4. Cluster proportions.**

**(A)** Bar plot showing the proportion of cells split by original identity. **(B)** UMAP showing the proportion of cells within each cluster (*res = 0.8*). Samples LD24-Int, LD26-Int, and LD28-Int are the "social interaction", while LD21-ctrl, LD22-ctrl, and LD31-ctrl are the "social observer".

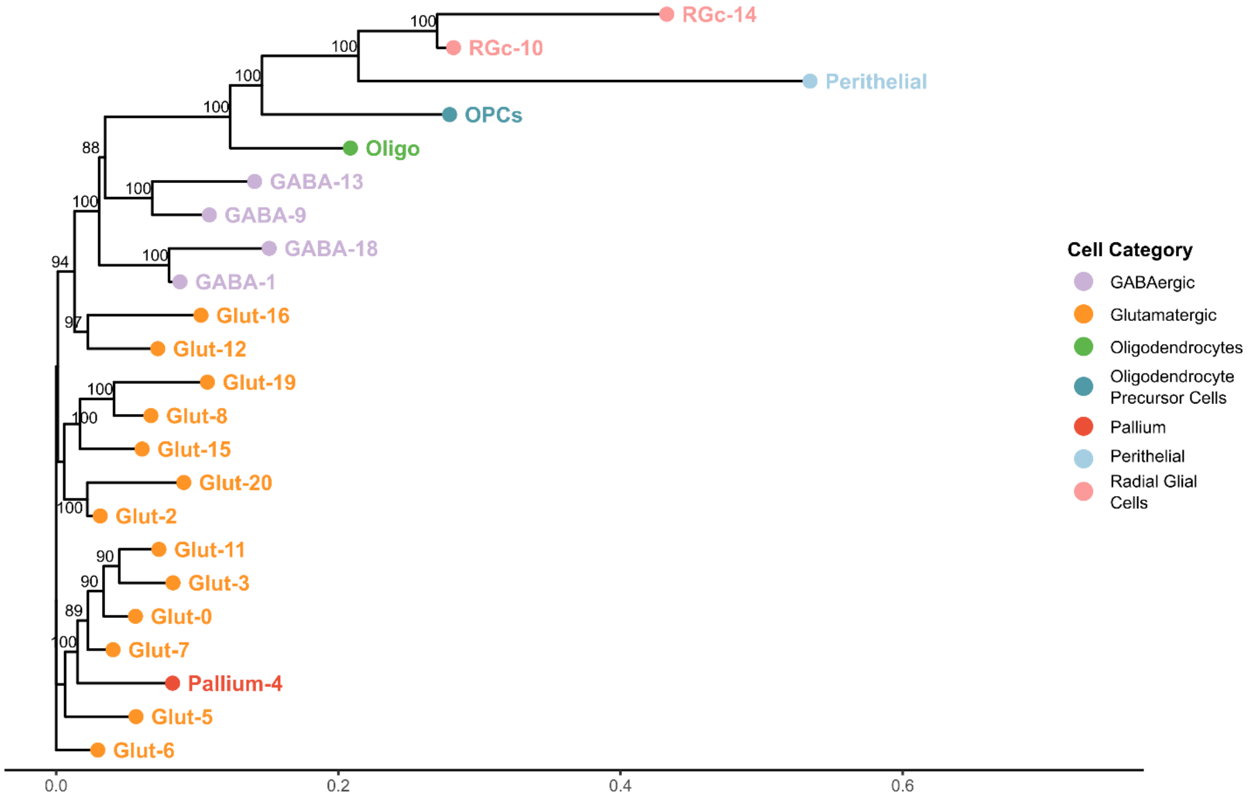


**Fig. S5. Cell type tree for the *L. dimidiatus* snRNA-seq dataset.** Unbiased cell type tree reconstructed using all expressed genes and 10,000 bootstraps.

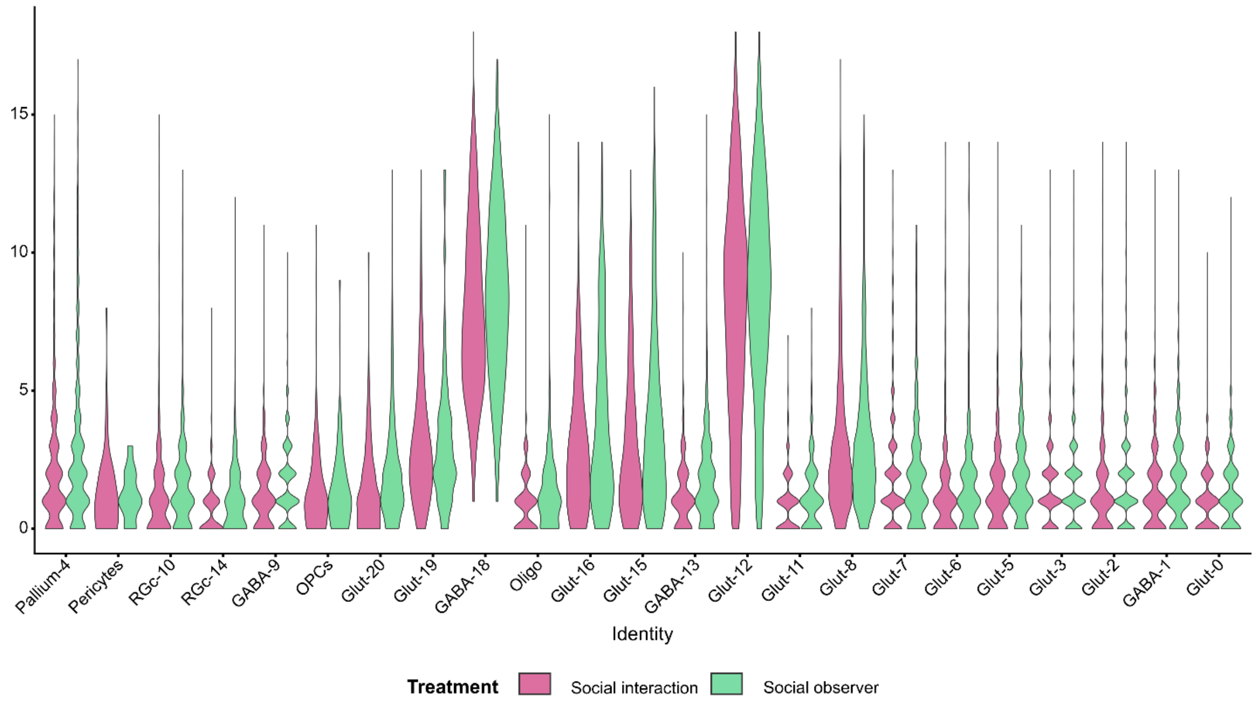


**Fig. S6. Violin plot of the IEGs-like distribution across cell clusters and treatment.** Co-expressed genes (IEGs-like) with *c-fos*, *egr1*, and *npas4* across cell clusters and treatment. The full list of the IEGs-like genes identified is reported in Table S8.

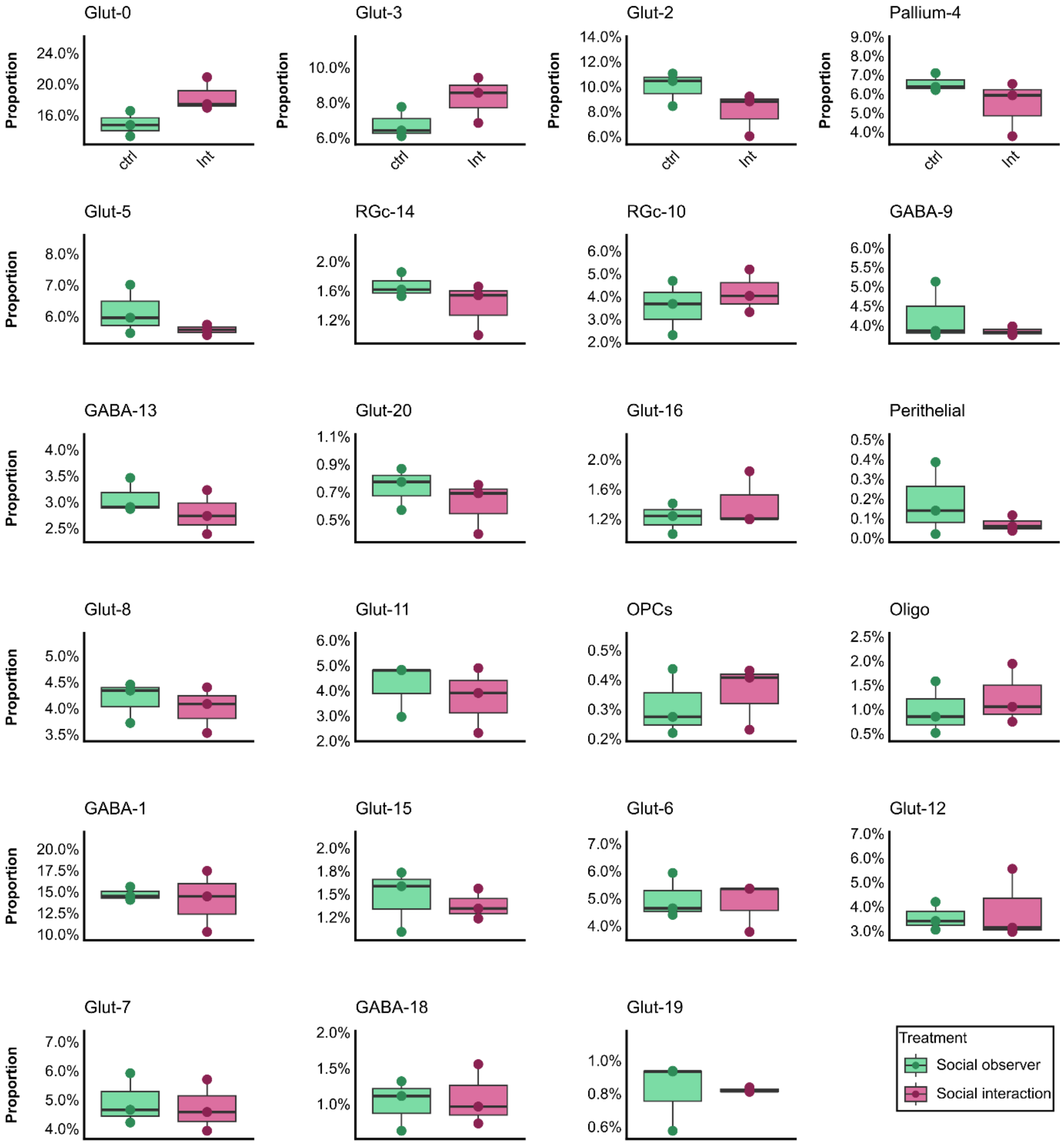


**Fig. S7. Boxplot with cell cluster proportion differences.** Adjusted p-values with the Benjamini-Hochberg correction (significant with $q < 0.05$). Results of the GLMM-Wald test are reported in Table S9A.

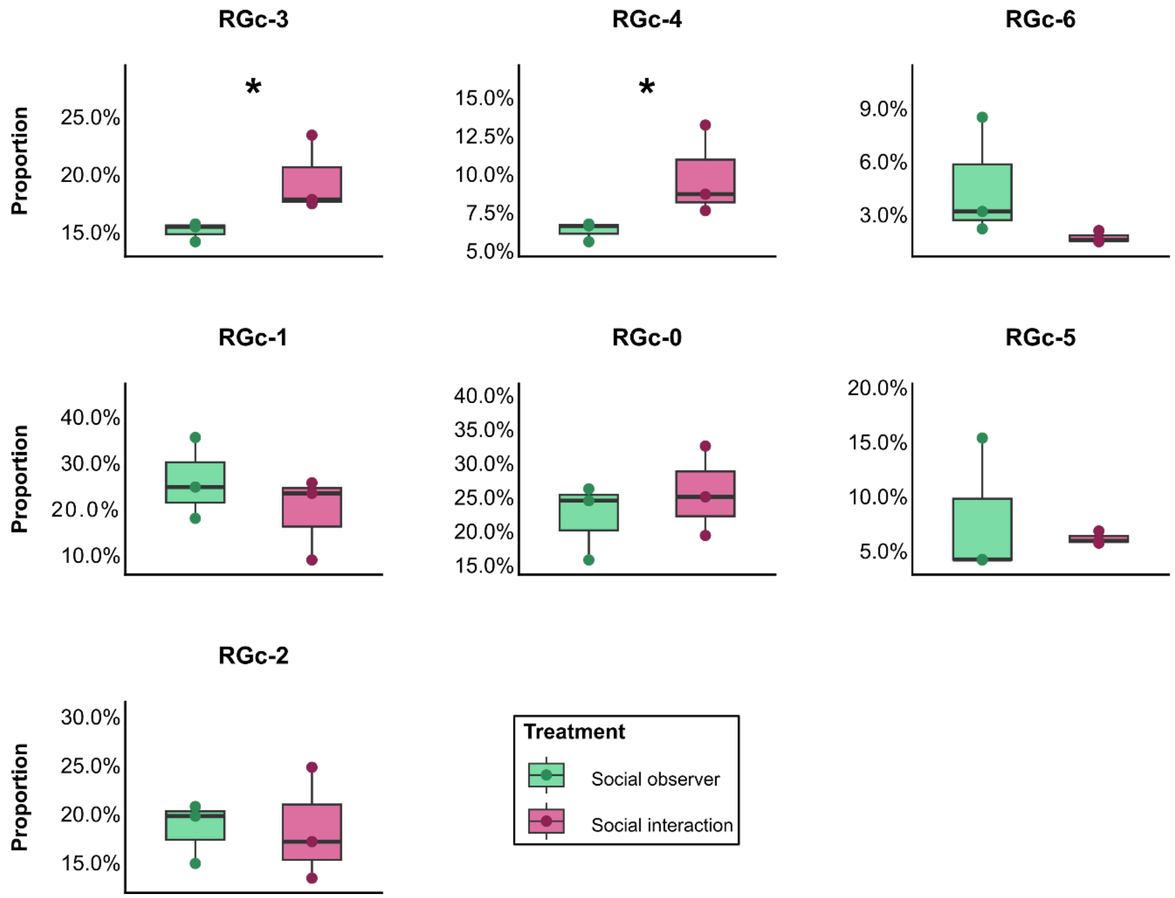


**Figure S8. Radial Glial (RGc) subclusters cell proportion.** Boxplot with cell proportion differences within the RGc subclusters (RGc-0 to RGc-6). Adjusted p-values with the Benjamini-Hochberg correction (significant with q < 0.05). Significant values indicate with "*". Results of the GLMM-Wald test are reported in Table S9B.

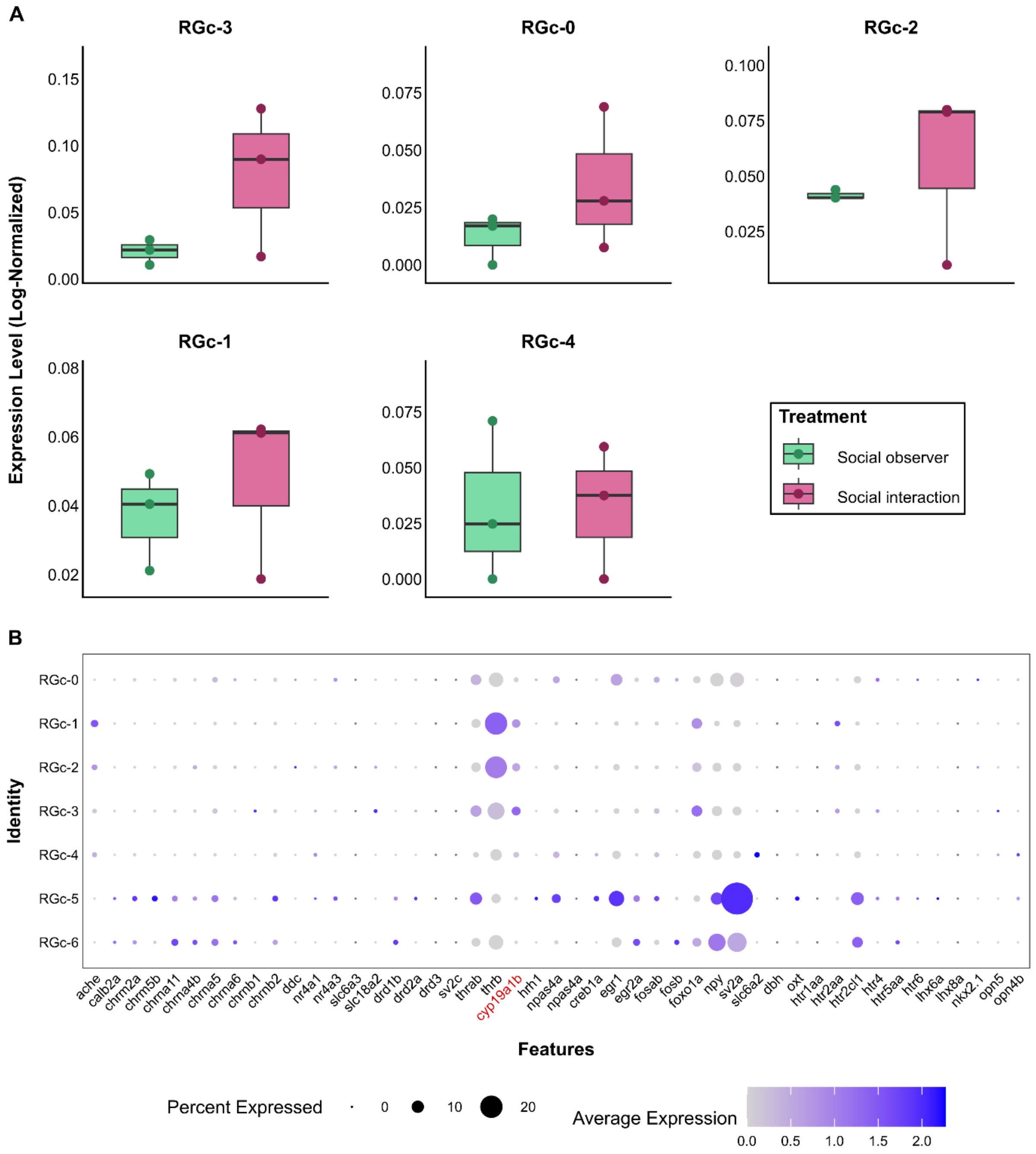


**Fig. S9. Radial Glial (RGc) subclusters. (A)** Boxplot with aromatase (*cyp19a1b*) expression among RGc subclusters. No significant differences identified (q < 0.05) **(B)** Dot plot showing the percentage of nuclei and the average expression profile of selected genes involved in social behaviour in *L. dimidiatus*. Aromatase gene (*cyp19a1b*) is highlighted in red.

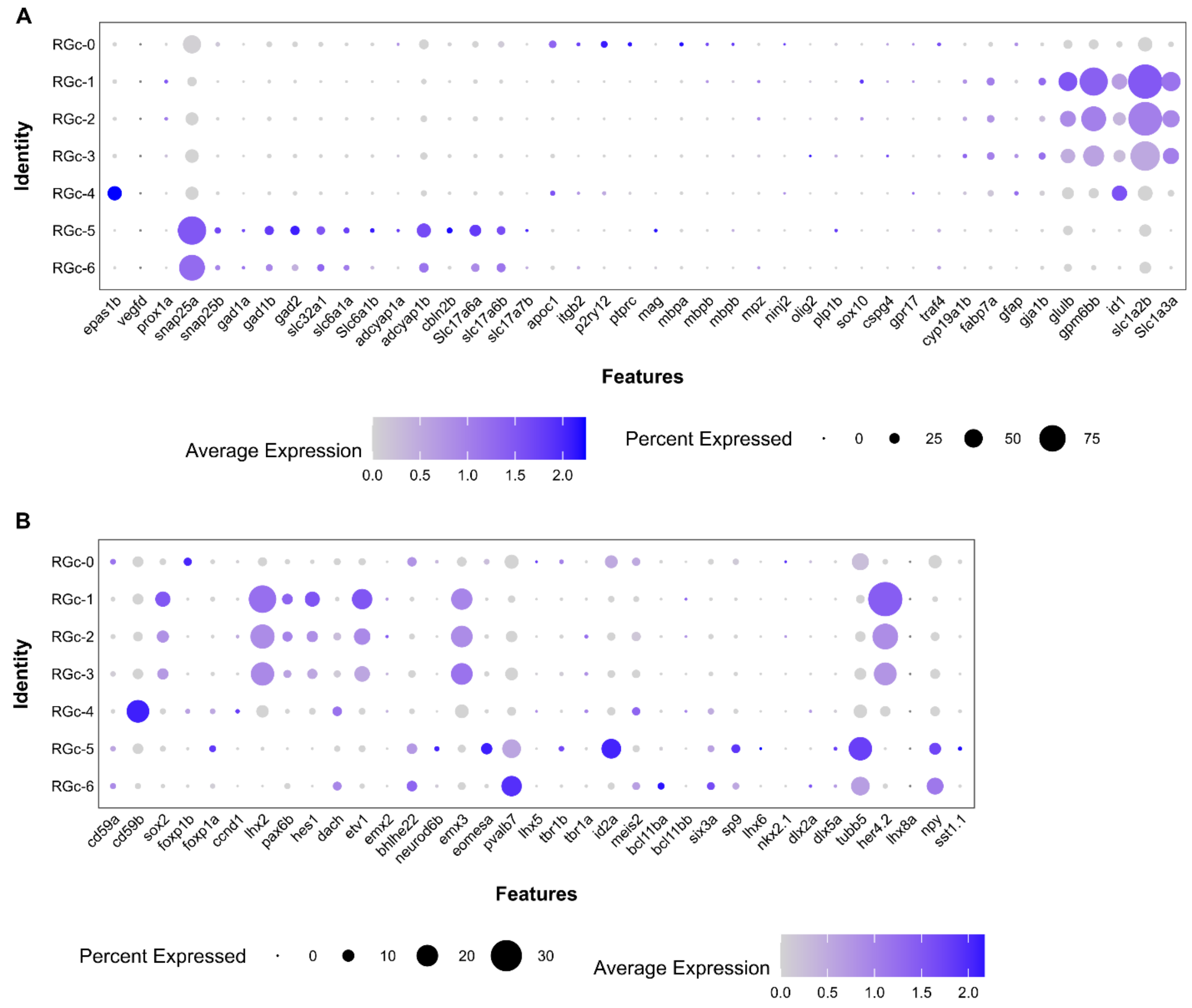


**Figure S10. Radial Glial (RGc) subclusters. (A)** Dotplot showing the percentage of nuclei and the average expression of canonical marker genes identified from the literature within RGc subclusters. **(B)** Dotplot showing the percentage of nuclei and the average expression of established neuroanatomically restricted priori genes of interest across the identified RGc subclusters.

### Supplementary Text

### Genome annotation pipeline

#### *Gene structure prediction and functional annotation pipeline*

The soft-masked genome assembly of *Labroides dimidiatus* (ASM3071049v1) was used as input for gene structure prediction by using BRAKER3 (v3.0.8) (*1*) inside Singularity container v3.8.5 (*2*). BRAKER3 employs an automatic pipeline that integrates transcript prediction, homolog evidence-based prediction and *ab initio* prediction to enhance the annotation accuracy. Brain specific RNA sequencing data from three different brain regions (fore-, mid-, hindbrain) and five more specific brain areas (including optic tectum, telencephalon, cerebellum, diencephalon, brain stem) were used to improve the annotation quality (for a total of 440 libraries, including data available under the NCBI BioProject PRJNA1120171, PRJNA726349, and unpublished RNA sequencing data). Raw sequencing data were first trimmed to remove adapters and poor-quality reads using Trim Galore! v0.6.10 (*3*) with the installed Cutadapt v4.5 *(--illumina -q 30, --length 50*). Then, the filtered data was aligned with STAR v2.7.11b with the "two pass mode" *(--outSAMstrandField intronMotif*) to improve the splice junction annotation (*4*, *5*). The resulting bam files were used as input to BRAKER3 *(--busco_lineage Actinopterygii_odb10, --species=Labroidesdimidiatus*) along with the vertebrata OrthoDB v12 partition (*6*), including specifically the Actinopterygii protein database (odb12.2025-07-01), and the protein sequences from *Danio rerio* (GCF_049306965.1) and *Halichoeres trimaculatus* (GCF_033239565.1). The obtained final high-confidence gene set included 23,173 protein-coding genes.

The quality of genome annotation was assessed using BUSCO v6.0.0 (*7*) with the lineage dataset "actinopterygii_odb12". We identified 97.5% of complete BUSCOs with 65.4% complete and single-copy BUSCOs, and only 0.6% fragmented and 1.9% missing BUSCOs (Table 1).

The predicted protein model was functionally annotated by aligning it against UniProt with InterProScan v5.75-106 (*8*) (*-goterms, -iprlookup*) and eggNOG mapper v2.1.12 (*9*) using diamond v2.0.11 (*--score 60, --pident 40, --query_cover 20, tax_scope 7742, --go_evidence all*) (*10*). In summary, 88.8% of the predicted genes were functionally annotated and the final *Labroides dimidiatus* annotation file can be retrieved at "https://www.schunterlab.com/resources".

**Table 1. BUSCOs results.** Quality of *Labroides dimidiatus* genome annotation assessed with BUSCO.

| BUSCO | |
|---|---|
| C:97.5%[S:65.4%,D:32.1%],F:0.6%,M:1.9%,n:7207 | |
| 7025 | Complete BUSCOs (C) |
| 4714 | Complete and single-copy BUSCOs (S) |
| 2311 | Complete and duplicated BUSCOs (D) |
| 44 | Fragmented BUSCOs (F) |
| 138 | Missing BUSCOs (M) |
| 7207 | Total BUSCO groups searched |

***Mitochondrial gene annotation***

The annotation for mitochondrial-specific genes present in the mitochondrial chromosome (CM061204.1) was carried out using MitoAnnotator programme (*11*) using default parameters. A total of 13 genes, 2 rRNAs and 22 tRNAs (Fig. 1) were included in the final *Labroides dimidiatus* GTF.

***3' untranslated regions (UTRs) annotation***

To improve the genome annotation quality, peaks2utr tool (*12*) was used to annotate 3' untranslated regions (UTRs). Peaks2utr call broad "peaks" of significant read coverage in one bam input file and uses those peaks to annotate novel 3' UTRs. To obtain a more comprehensive picture and stronger evidence of 3' UTRs presence, two bam files per each brain region from the sequencing libraries previously used to generate the genome annotation were randomly chosen and merged using Samtools v1.22.1 (*13*). One random sample from the current single-nuclei RNA sequencing experiment (LD22 -social_obs) was also included to generate the final input bam file. The resulting GTF was then used as input for Cell Ranger v6.0.1 pipeline.

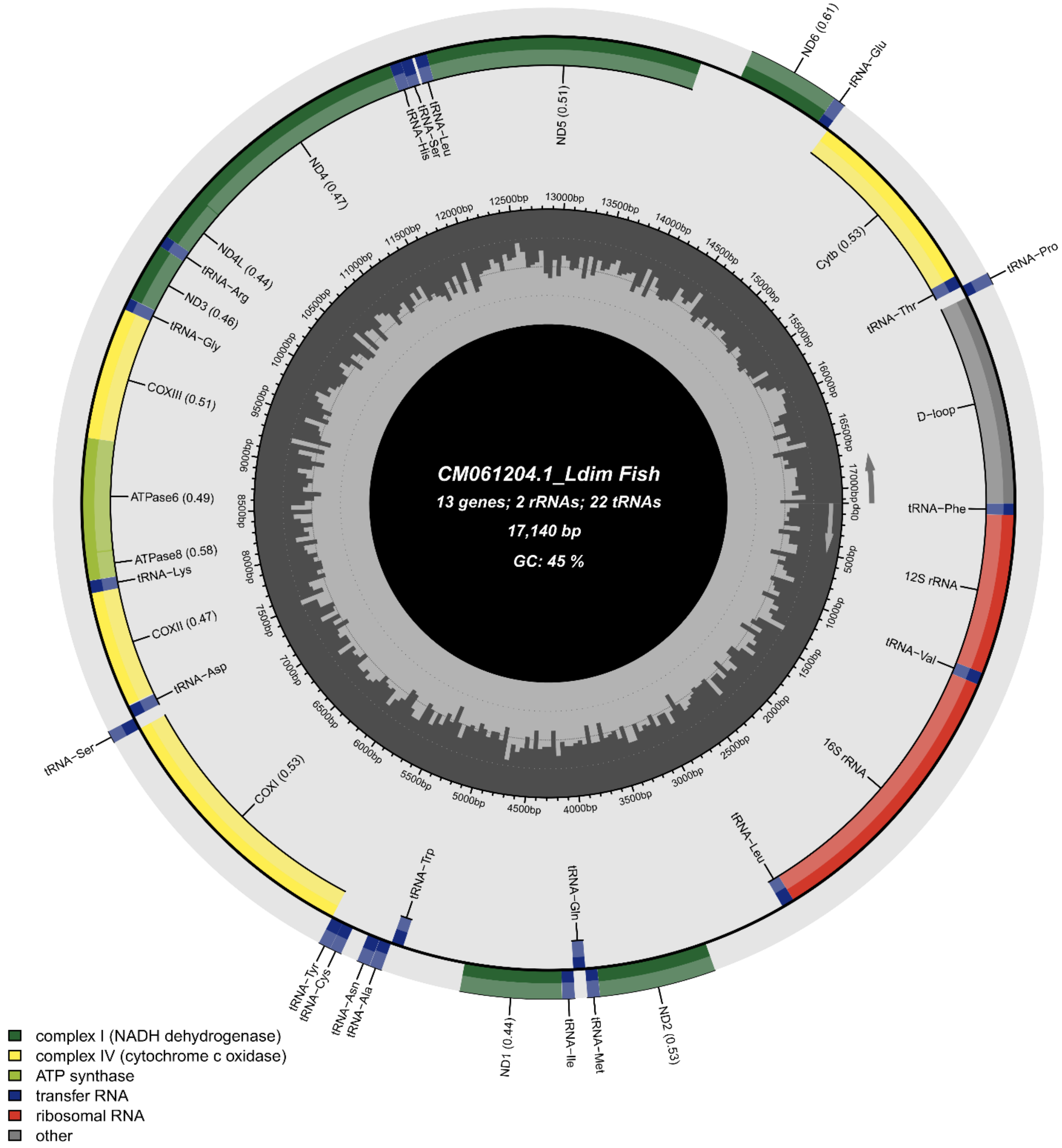


**Fig. 1. Mitochondrial-specific genes present in the mitochondrial chromosome (CM061204.1) of *L. dimidiatus*.** Ribosomal RNA (rRNAs), transfer (tRNAs).
